# A functional SNP regulates E-cadherin expression by dynamically remodeling the 3D structure of a promoter-associated non-coding RNA transcript

**DOI:** 10.1101/2021.10.14.464445

**Authors:** Shrikant Sharma, Gabriele Varani

## Abstract

Transcription of E-cadherin, a tumor suppressor which plays critical roles in cell adhesion and the epithelial-mesenchymal transition, is regulated by a promoter-associated non-coding transcript. This RNA includes a functional C/A single nucleotide polymorphism (SNP rs16260). The A-allele is linked to decreased transcriptional activity and increased prostate cancer risk. This single nucleotide change affects recruitment of an iso-miRNA and epigenetic enzymes to regulate the promoter, yet it is distant from the isomiR-binding site in both primary sequence and secondary structure, raising the question of how regulation occurs. Here we report the 3D NMR structure of the domain of 90 nucleotides within the paRNA which includes the SNP and the isomiR-binding site. We show that the A->C mutation alters the locally dynamic structure of the paRNA, revealing that the mutation regulates the E-cadherin promoter through its effect on RNA structure.

## Introduction

Classical cadherins, such as E-cadherin^1^, are transmembrane glycoprotein components of adherens junctions which promote intercellular communication^2,3^. The E-cadherin gene^4^ (CDH1) generates a 120 kDa protein^5^ whose cytoplasmic domain links various catenins to the actin cytoskeleton and facilitates downstream signaling through multiple pathways, including Wnt and TGF-β^6,7^. Mis-function of E-cadherin is linked to invasiveness and advanced tumor stage in many epithelial cancers^8,9^. In fact, reduced E-cadherin expression is a hallmark of the epithelial-mesenchymal transition (EMT)^10^, and inhibition of E-cadherin function provokes separation and invasion of cancer cells^11^. Thus, E-cadherin is a tumor suppressor whose deregulation promotes carcinogenesis^12^.

Hypermethylation of the CDH1 promoter has been observed in human breast, prostate, and hepatocellular tumors carrying a wildtype CDH1 gene, leading to reduced expression of E-cadherin^13,14^. In addition, a C->A (SNP rs16260) polymorphism at position -160 from the transcriptional start site decreases the activity of the CDH1 promoter by about 70% and is linked to increased risk for prostate cancer^15^. The C allele more robustly recruits transcription factors compared to the A allele^15^. Mechanistically, it was demonstrated that silencing of the CDH1 promoter requires formation of a microRNA (miRNA)-guided Argonaute 1 (AGO1) complex on an independently transcribed sense promoter-associated transcript, which encompasses the SNP and recruits the SUV39H1 methyltransferase to affect chromatin modifications^16^ (Fig. 1A). It was also demonstrated that the SNP-160 (C/A) influences the ability of isomiR-4534 and AGO1 to interact with the paRNA^16^. However, it remains unclear how the polymorphism affects transcriptional regulation, since it does not overlap with the isomiR binding site in sequence or secondary structure. The A and C variants differed substantially in the pattern of SHAPE reactivity at and near SNP rs16260, revealing different secondary structures for the two polymorphic transcripts^16^, but how the secondary structure changes is unclear too, because the A is single stranded in the proposed secondary structure model.

**Fig. 1.**
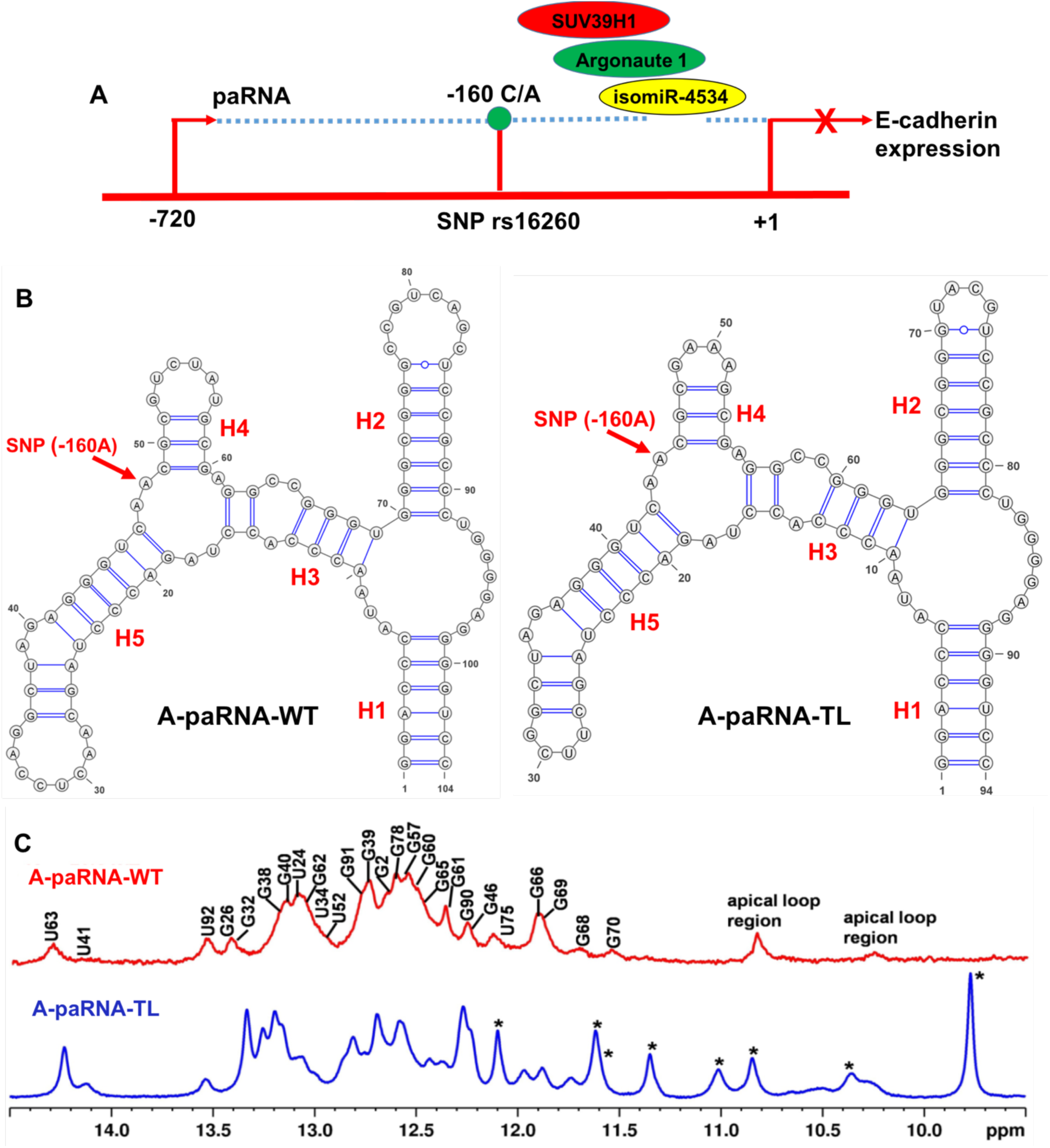
A) Schematic diagram of the impact of SNP rs16260 on transcriptional regulation of E-cadherin. B) Secondary structure of the domain of the CDH1 paRNA that regulates E-cadherin transcription, named A-paRNA, and of its variant A-paRNA-TL, where tetraloops were introduced to stabilize the RNA structure and reduce aggregation which caused poor spectral quality. C) Comparison of the 1D imino ^1^H NMR spectra of the two RNAs, with assignments for the A-paRNA-TL, obtained as presented in the text, mapped onto the spectrum of the sequence with wild type loops. The spectra are very similar, indicative of conserved structures; (*) identify resonances originating from the UUCG, GAAA and UACG tetraloops, within the A-paRNA-TL construct.

We report the three-dimensional structure of the 90 nucleotides domain within the promoter associated RNA containing the A-SNP, determined using NMR spectroscopy. We show that the paRNA folds into a well-defined three-dimensional structure, and that the A->C mutation unfolds a partially dynamic three-way junction where the SNP is located. Propagation of this rearrangement to a neighboring dynamic internal loop extends the structural reorganization to the helix that links the three-way junction to the isomiR binding site. Thus, we suggest a new mechanism for regulation of ncRNA function which emphasizes the functional role of RNA structure and its modulation by genetic variation. Since changing the structure of ncRNAs at polymorphic sites and through somatic mutations can affect the epigenetic landscape of human disease^14-16^, including cancer, the potential for therapeutic targeting should be considered.

## Results

### Secondary structure of the E-cadherin transcript with the -160A allele (A-paRNA)

We recorded NMR spectra for the A-paRNA, corresponding to the domain of the sense transcript with the -160A allele. The 1D spectrum contain many NH peaks in the region where base pairs can be directly monitored (11-14 ppm), revealing a well-folded structure (Fig. 1). However, the relatively poor quality of the spectra, as reflected as well in broad and overlapped cross-peaks in 2D non-exchangeable protons (Fig. S1), would prevent structure determination. In fact, even for exchangeable peaks, below 15 °C, the spectra become broadened to the point of being nearly useless. We encountered this problem with several other RNAs studied in the laboratory, including the CssA thermometer^17^ and the stem loop within the c-JUN 5′ UTR recognized by eIF3 during specialized translation initiation^18^. As was done in those projects, we introduced tetraloops in the A-paRNA sequence, in place of three existing loops, to generate A-paRNA-TL, as shown in Fig. 1B, based on the SHAPE-generated secondary structure^16^. This is not unlike what is often done in RNA x-ray crystallography, where protein binding sites and tertiary contacts are engineered to stabilize crystal contacts and facilitate crystallization. We reasoned that introducing tetraloops would stabilize the secondary structure and reduce aggregation; indeed, the spectral quality improved considerably (Fig. 1C).

To examine whether introduction of the tetraloops altered the RNA structure, we mapped the NH assignments for A-paRNA-TL (with tetraloops), obtained as described below, onto the A-paRNA (wild type loops) spectra. An overlay of the imino region spectra for the two RNAs show many overlapping peaks, indicative of very similar secondary structures for both constructs (Fig. 1C). Thus, introducing tetraloops in the A-paRNA sequence does not change its secondary structure but stabilizes it and reduces aggregation, making high resolution NMR studies possible.

The sharp and well-resolved H2 protons allowed us to identify U-A base pairs, while G-C base pairs were identified from the strong cross peaks between GH1 and the pair of Cytidine amino protons. Wobble base pairs were identified from the characteristically strong cross peaks between GNH1 and UNH3 within the non-canonical base-pairing range (12-10 ppm). Based on this attribution, and the canonical pattern of sequential NOEs involving imino protons in helical regions, we assigned all slowly exchanging NH chemical shifts for the A-paRNA-TL and for a shorter variant lacking the terminal stem-loop (called A-paRNA-TL-tr, Fig. 2), designed to isolate the 3-way junction where the SNP is located, the two stem-loops emanating from it (H4 and H5), as well as helix H3, but missing the larger 3-way junction near the boundary of the domain formed by helices H1, H2 and H3. All assignments were consistent with the predicted secondary structure, and all base pairs predicted from the SHAPE analysis were observed, except for nucleotides that are paired but at the end of helices; imino protons for unpaired nucleotides were instead not observed; overall, the results are fully consistent with the secondary structure generated from the SHAPE analysis.

**Fig. 2.**
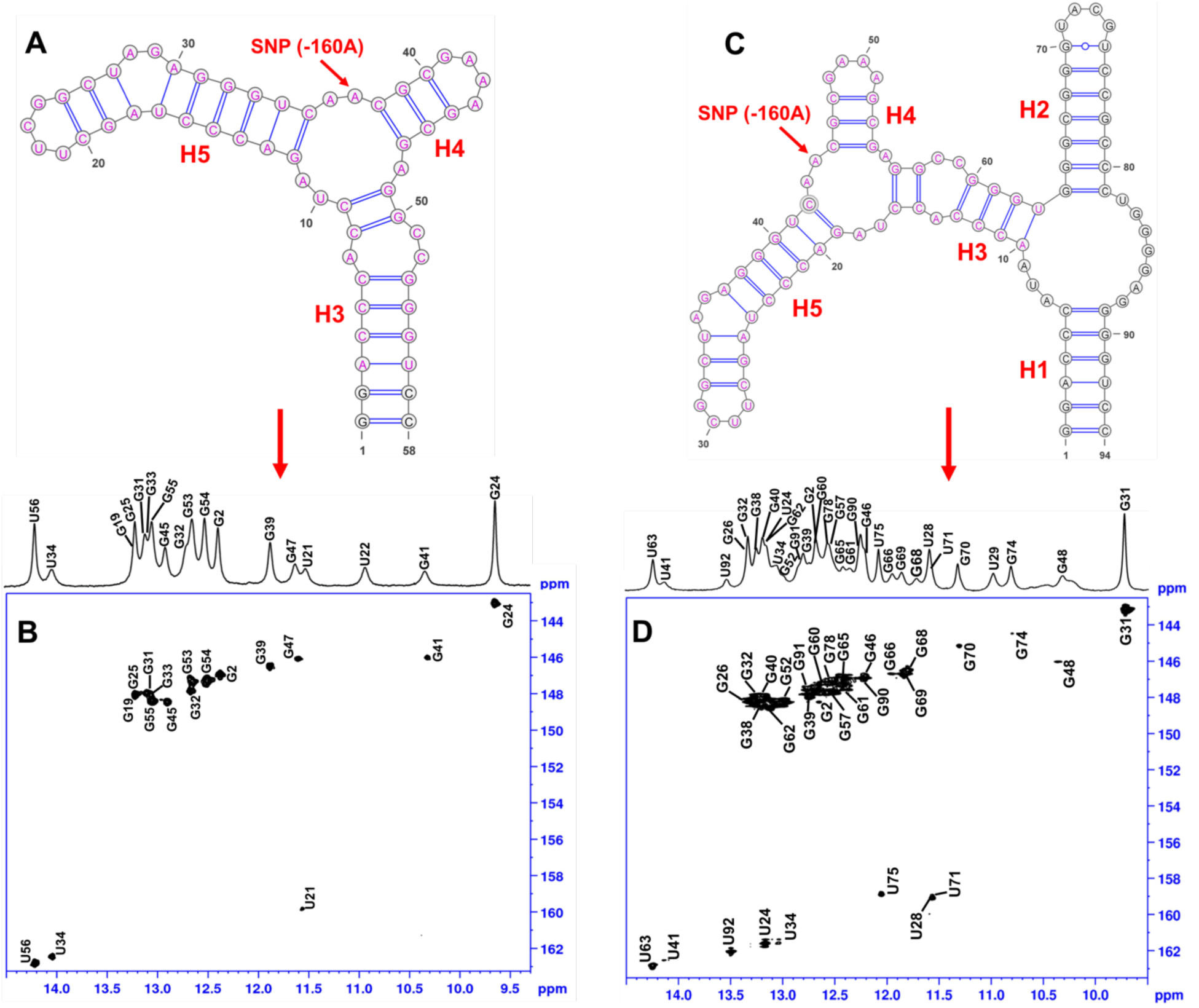
A) Secondary structure of A-paRNA-TL-tr, which was designed to isolate the three-way junction where the SNP is located, as established by NMR; B) 2D-^15^N-HSQC spectra of the same RNA construct, with NMR assignments. C) Secondary structure of A-paRNA-TL, as established by NMR; D) 2D-^15^N-HSQC spectra of the same RNA construct, with NMR assignments.

In order to investigate whether the C/A polymorphism would alter the secondary structure, we prepared and purified the C-paRNA construct with the -160C allele in place of A (Fig S2). As before, we introduced tetraloop, consistent with the secondary structure established from SHAPE, to improve spectral quality and obtain complete assignments. An overlay of the 1D imino region for the A-paRNA-TL (−160A allele) and C-paRNA-TL (−160C) (Fig. S2) show that the NMR spectra of A and C variants differ substantially, reflecting changes in the secondary structure due to the SNP. To confirm this conclusion, we assigned the NHs of the C-paRNA-TL (−160C) as well, and established its secondary structure based on the NH assignments (Fig. S2). Based on this analysis, we conclude that the region of the two RNAs that comprise the isomiR binding site (helix H2 in Fig. 1) adopts the same secondary structure regardless of the A- or C-allele at position -160. Furthermore, the longer stem-loop emanating from the three-way junction where the SNP is located (helix H5 in Fig. 1) also retains the same secondary structure, as does helix H1 that defines the boundaries of the domain. However, the single nucleotide A-C change at position -160 within the three-way junction region alters the structure of the junction and results in a new secondary structure for the junction itself, of helix H4 as well as helix H3 that links the SNP to the domain that recruits the isomiR. As a consequence of the rearrangement of helix H3, the other three-way junction is also remodeled, so that helix H2 now emanates from a larger multi-helix junction. It is conceivable that this new structure would change the thermodynamic or kinetic stability of helix H2, allowing differential access by isomiR-4534 which initiates assembly of SUV39H1.

In summary, the analysis of the NMR spectra of the paRNA domains containing either A- and C-alleles demonstrates that both RNAs are well-folded, consistent with the SHAPE analysis^16^, and that the secondary structures of the transcripts of the two polymorphic sequences differ in the region linking the SNP to the isomiR binding site. To understand which structural features of the RNA are responsible for the conformational change, we set up to determine the three-dimensional structure of the A-paRNA-TL construct.

### ‘Divide-and-conquer’ allows nearly complete spectral assignments and the collection of a large NOE dataset

The quality of the NMR spectra of the A-paRNA-TL construct is very high considering the size of the RNA, but it remains very challenging to obtain complete spectral assignments and collect a large number of experimental NOEs, due to fast relaxation leading to peak broadening, as well as very crowded NMR spectra. 3D and higher dimensionality experiments based on ^13^C editing are of limited use due to fast relaxation, as is segmental labeling that would reduce spectral overlap but not the broad lines. Limited experimental NMR data (the few NOEs from slowly relaxing protons like AH2 obtained by extensive per-deuteration and selective carbon labeling) have been combined with RDC’s and SAXS to generate 3D structures, but these approaches rely extensively on modeling assumptions to overcome the paucity of direct experimental observations^19-22^. Unsurprisingly, the quality of NMR structures of RNA improves significantly when the density of experimental NOEs is high^17,18,23^.

To address these limitations, we used the proven ‘divide and conquer’ approach to complete chemical shift assignments and obtain a large number of NMR constraints for structure determination, starting with smaller independently folded sub-domains. This approach, illustrated in Fig. 2 and Fig. S3, is grounded in the modular nature of RNA structure; in analogy to multidomain proteins, RNA secondary structure domains fold independently and tertiary interactions stabilize but only seldom rearrange the secondary structure of the RNA^24-26^. Thus, several smaller constructs of varying length were generated (18-58 nts; Fig. S3). When common and overlapping base pairs are superposed, these constructs combine to generate the complete A-paRNA-TL (Figs. S3 and S4). The larger fragments (called A-paRNA-TL-tr and A-paRNA-TL-tr-1) comprise the 3-way junction formed by helices H3, H4 and H5 (Fig. 1; Fig. 2) where the A/C SNP is located, and isolate it within smaller RNAs to reduce spectral overlap and allow closer investigation of its folding; the stem-loop derived from it (A-paRNA-TL-tr-2) isolates the longer stem-loop emerging from the 3-way junction, helix H5; finally, A-paRNA-2 isolates the isomiR-4534 binding site, helix H2.

When we examined the isolated sub-domains, we observed all the imino peaks in the 1D (Fig. S5) and 2D NMR spectra expected from the secondary structures shown in Figs. 1, 2 and S3; relevant illustrative spectra are shown in Figs. S6-S10. Overlaying the 1D and 2D-NOESY NMR spectra (with and without per-deuteration) of the smaller subdomains on the full A-paRNA-TL spectra revealed a highly transferable pattern of chemical shifts and NOESY cross-peaks for both the exchangeable and non-exchangeable protons (Figs. S4-S5 and S11), demonstrating that the structure found in the complete A-paRNA is retained in each fragment. Once the secondary structure of each fragment was verified, we recorded 2D and 3D-NOESY spectra at different mixing times for both exchangeable and non-exchangeable protons to obtain nearly complete and unambiguous chemical shift assignments. To achieve further simplification of the spectra, we also prepared per-deuterated RNA samples for the three larger fragments, A-paRNA-tr-1, A-paRNA-tr-2 and A-paRNA-tr (representative sections of the spectra are shown in Figs. S12-S14) and compared the resulting NMR spectra with the spectra of the perdeuterated A-paRNA-TL (Figs. S11 and S15). Briefly, imino protons for A-paRNA-TL establish its secondary structure and were initiated with the U28-G32, G70-U75 and U71-G74 wobble base pairs identified by strong cross peaks between GH1 and UH3 within the non-canonical base-pairing range (12-10 ppm) (Fig. S10). A series of D_2_O 2D NOESY experiments with mixing times between 100 and 300 ms were used to assign the non-exchangeable protons. The overlap in the NMR ribose spectra was resolved through deuteration of the H3′, H4′ and H5′/ 5′′ positions in the sugar and the H5 positions of C/U, which simplified the spectra and sharpened the linewidth by reducing dipolar relaxation (Fig. S16). Nearly complete spectral assignments were obtained in this manner. Altogether, with the aid of deuteration and 3D ^13^C-edited NOESY spectra, we assigned 91.5% of resonances for A-paRNA-TL (significant exceptions are the H3′, H4′, H5′ and H5′′ peaks for residues U8, U24, A25, A35, G36, A55, C58 and U82, which we were unable to assign). Based on these assignments, we then generated a comprehensive list of distance restraints for each of the individual fragments, which were merged to generate a restrain table for the A-paRNA-TL (Table 1). These restraints were used in the structure calculation of A-paRNA-TL.

**Table 1.**
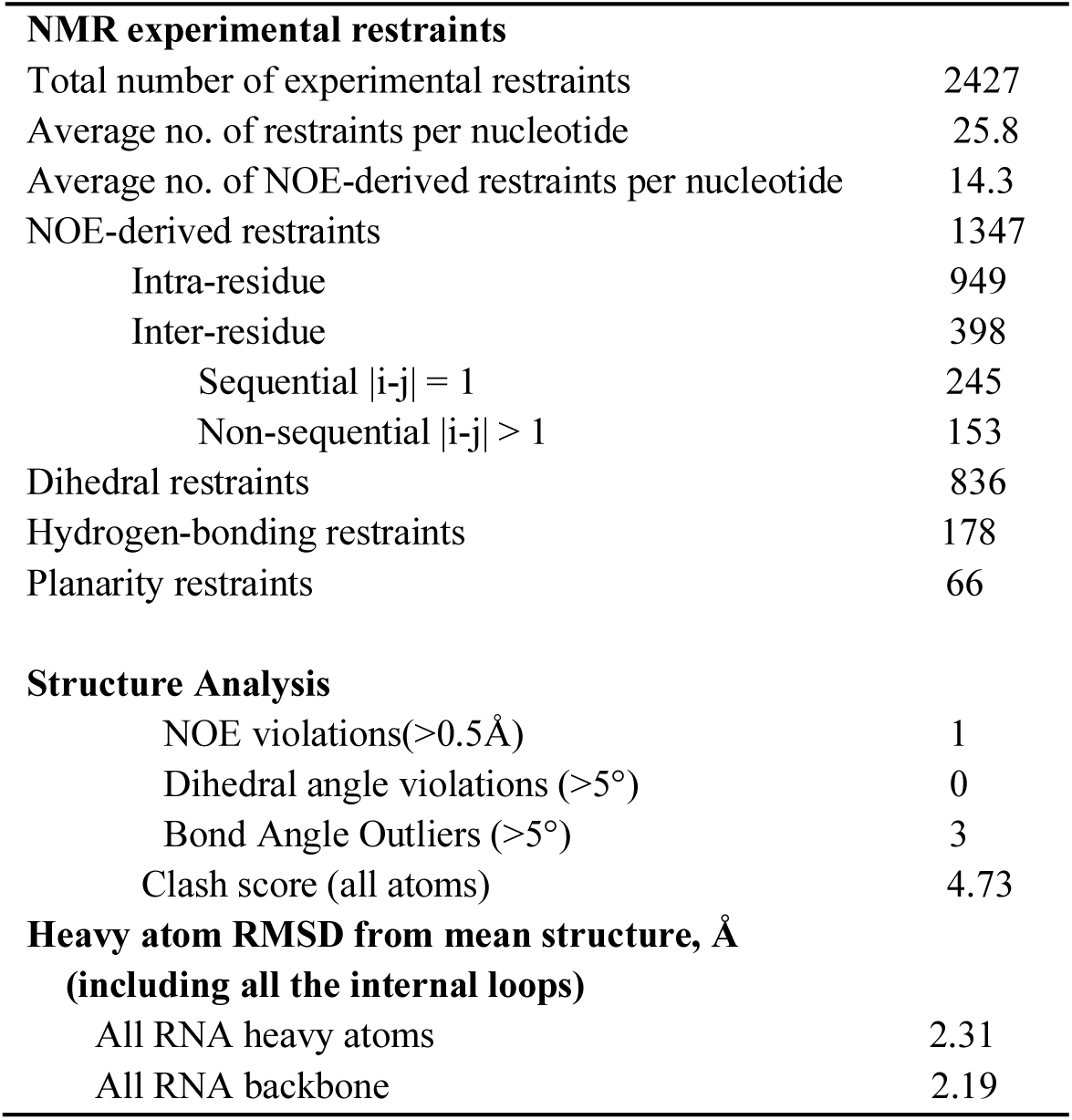
NMR and Structural Statistics for Structure Determination of A-paRNA-TL.

### Quality of the structure

The ‘divide and conquer’ approach allowed us to obtain a much larger number of constraints for structure determination than would have been possible if we just studied the complete RNA, whose large and asymmetric structure leads to rapid relaxation and broadening of the NMR signal. These physical limitations cannot be overcome by selective isotopic labeling. We generated the restraint set (Table 1) by combining NOE-derived distance and torsion angle restraints from each fragment and the structure was then calculated using a well-tested simulated annealing protocol (as discussed in methods section) within NIH-XPLOR^27^. A final ensemble of the 10 lowest energy structures (Fig. 3A) from a refined set of 150 structures was chosen for analysis. The overall RMSD (including all internal loops and three-way junctions) for backbone and heavy atoms were 2.19 and 2.31 Å, respectively. We attributed this relatively low RMSD value to the high number of NOE restraints per nucleotide obtained by building the final restraint sets from individual subdomains that overlap to generate the full RNA.

**Fig. 3.**
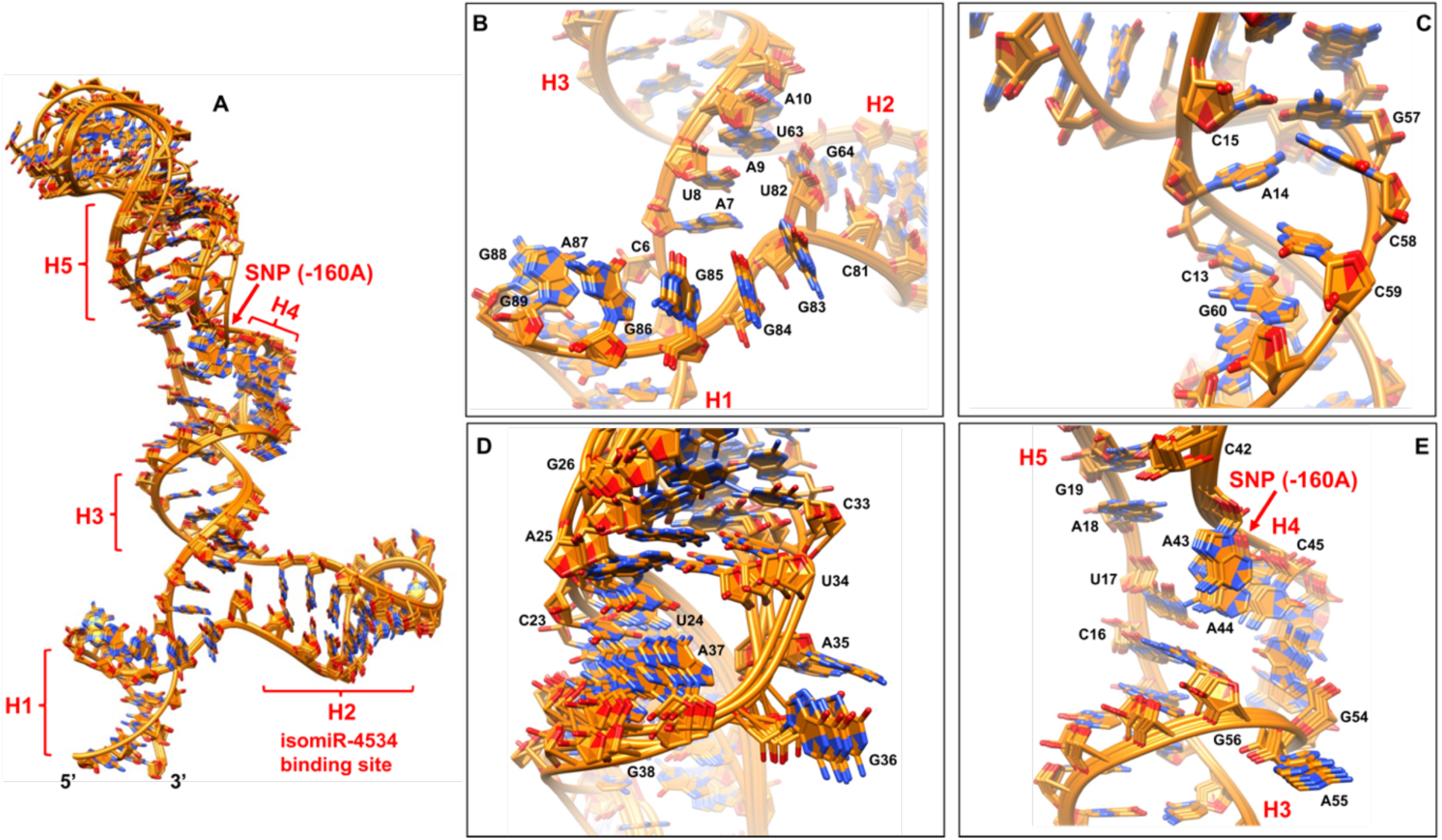
A) 10 lowest-energy structures superposed to generate the structural ensemble of A-paRNA-TL. B) Close up view of the 3-way junction from which the isomiR binding site emanates, formed by helices H1, H2 and H3. The unpaired bases G83-G89 stack on top to each other, while U82 is bulged out and U8 is inserted between residues A7 and A9 but not bulged out; as a result, helices H1 and H2 are coaxially stacked while helix H3 is in an orthogonal orientation; C) Close up view of the 13-15; 57-60 internal loop within helix H3; the unpaired A14 nucleotide is inserted between the G60-C13 and G57-C15 base pairs; the two bases on the opposite strand, C58 and C59, are displaced from the helix to accommodate imperfect stacking of the G60-C13 and G57-C15 base pairs; D) In the internal loop within helix H5, A35 and G36 are transiently bulged out to accommodate the U34-A25 and U24-A37 base pairs; this region of the structure displays relatively high flexibility, resulting in less precisely defined local structure; E) close up view of the three-way junction formed by helices H3, H4 and H5, where the SNP is located; A43 and A44 stack on top of each other and A55 is bulged out, while U17 and A18 are sandwiched between the G56-C16 and G19-C42 base pairs; as a result, helices H3 and H5 coaxially stack while the third helix H4 is in an orthogonal orientation.

Superposition of individual helical sections of the RNA are all below 1.00 Å; for example, helix H1 has RMSD=0.95 Å and helix H2 converges to a RMSD=0.90 Å (Fig. 3A). One exception is represented by the flexible tip of helix H5 (Figs. 2C and 3D), near the internal loop formed by nts 24-25/34-37, whose bases are transiently in a solvent exposed conformation that affect the neighboring base pairs as well. This region converged more poorly (Fig. 3D) due to the paucity of NOEs and shows a higher local RMSD value of 2.95 Å. Likewise, bases located within the SNP three-way junction formed by helices H3, H4 and H5 (e.g. nucleotides 16-19, 42-45 and 54-56), also converged less well (Fig. 2C and Fig. 3E) due to the paucity of NOEs between loop residues, leading to a locally higher RMSD =1.98 Å. In contrast, the internal loop involving nucleotides 13-15 and 57-60, located in helix H3 connecting the two three-way junctions (Figs. 2C and 3C) shows a low RMSD=1.19 Å, yet it remains conformationally dynamics, as evidenced by mixed sugar conformations, as discussed in the next paragraph. Likewise, the purine rich three-way junction formed by helices H1, H2 and H3 and involving nucleotides 6-10, 63-64 and 81-89, from which the isomiR-binding stem-loop H2 emanates (closed by the U63-A10, G64-C81 and G89-C6 base pairs, Figs. 2C and 3B) superposed very well (purines G83, G84, G85, G86, A87, G88 and G89 all stack on top to each other) and shows a low local RMSD=0.92 Å.

All nucleotides within the A-paRNA-TL construct show 3′-endo sugar pucker conformation with relatively few exceptions (we ignore in this discussion residues within the tetraloops whose properties have been described exhaustively in the past). These are A14 (within the 13-15 and 57-60 internal loop in helix H3); A35 and G36 (in the 24-25/34-37 internal loop within helix H5); A55 (within the SNP three-way junction), as well as U82 within the other three-way junction. All these residues exhibited strong H2′-H1′ and H3′-H1′ TOCSY correlations, suggesting 2′-endo sugar pucker conformation. In addition, C58 and C59 (within the 13-15 and 57-60 internal loop in helix H3, Figs. 2C and 3C) as well as U17 (part of the SNP three-way junction (Figs. 2C and 3E), exhibit mixed 2′-endo/3′-endo sugar pucker conformation, based on relatively strong H1′ to H2′, H3′, and H4′ cross-peaks, and also display deviations from A-form helical pattern in the backbone, consistent with the presence of local conformational dynamics. This dynamic is reflected in higher local uncertainty in the structure and higher RMSD, as discussed in the previous paragraph. As discussed next, the conformational flexibility of the three-way junction where the SNP is located as well as internal loop in helix H3 is likely to be functionally significant.

### 3D structure of the CDH1 paRNA (−160A SNP)

The first feature of the structure, starting from its 5′- and 3′-ends that come together to form helix H1 that defines the boundaries of the domain, is the purine-rich three-way junction abutted by helices H1, H2 and H3 (Figs. 2C and 3B). No evidence of stable cross-strand base pairing was found based on the absence of NOEs between guanine NH2 or AH2 to the cross strand H1′, or the observation of slow exchanging NHs. However, the stretch of purines G83-G89 bases all stack on top to each other (Fig. 3B), as reflected in the characteristic sequential pattern of NOEs from H2′/ H1′ to H6/H8 (Fig. S17) observed in the highly deuterated sample; only residue U82 is bulged out. U8 is inserted between A7 and A9 but not bulged out, allowing for stacking interactions between A7 and A9 (Fig. 3B). As a result, two of the helices stack coaxially (H1 and H3), while the third helix (H2) is oriented more or less orthogonally to the coaxial stack (Fig. 3B). This topological arrangement matches the ‘family A’ topology of three-way junctions^28^, a very common tertiary motif in many three-way RNA junctions^28,29^.

The most important structural and functional feature of the paRNA is the three-way junction formed by helices H3, H4 and H5 (Fig. 2C and Fig. 3E), where the SNP is located. Within this junction, residues A43 and A44 stack on top of each other while A55 is bulged out (Fig. 3E); residues U17 and A18 are sandwiched between the G56-C16 and G19-C42 base pairs (Fig. 3E). In the NMR structure, residue A44, which coincides with the SNP-160A, is unpaired and stacked on A43 but does not form any cross strands interactions. Pairs of unpaired A’s are often found in 3-ways junctions and internal loops, for example in ribosomal RNAs^30,31^. These residues often orient and stabilize coaxially stacked helices at three-way junctions and are important for the formation of tertiary motifs in these RNAs^30,31^.

This ‘SNP-three-way’ junction shows similar topology and arrangements of helices as the first three-way junction, with two helices coaxially stacked (H3 and H5) and the third helix (H4) orthogonal to them (Fig 3E). However, this second three-way junction (Fig 3E) is less conformationally rigid than the first (Fig 3B; Fig 3E), with a local RMSD=1.98 Å. This conformational flexibility could facilitate the structural rearrangement that occurs when A-160 is changed to C. In the C-paRNA (−160C), the AA motif is changed to AC. When this mutation is introduced, the entire three-way junction unfolds in conjunction with helix H3 linking it to the larger three-way junction near the isomiR binding site. As a result, a much larger four-way junction forms (Fig. S2). This reorganization is most likely facilitated by the small internal loop within H3 just two base pairs away from the SNP junction, which introduces a structural distortion that destabilizes helix H3, which is conformationally dynamic, as revealed by the mixed sugar conformation observed for several residues.

In the internal loop within helix H3 involving nucleotides 13-15 and 57-60, A14 is inserted between the G60-C13 and G57-C15 base pairs (Fig. 3C) and the only deviation from A-form helical pattern is found in the C15 backbone angles, as required to accommodate A14. The two bases opposite, C58 and C59, however, are displaced from helical stacking and experience conformational exchange between 2′-endo and 3′-endo pucker, like U17 in the 3-way junction. The three unpaired bases A14, and especially C58 and C59, provide conformational flexibility to the internal loop and destabilize helix H3, which could facilitate the rearrangement of the entire helix when the A-C mutation occurs, which is necessary to remodel the secondary structure.

To test the hypothesis that the stability of the internal loop would affect the conformation of the paRNA domain, we synthesized two additional RNAs with wild type loops corresponding to either A and C-containing paRNA, truncated to helix H3 (Fig. S18). The CC motif (nts 58 and 59) is replaced with U in each of the two RNAs to create a perfectly base paired helix, containing either A- and C-allele at the neighboring 3-way junction. Comparison of the 1D imino ^1^H NMR spectra (Fig. S19) of the three RNAs shown in Fig. S18 are very similar, except for the appearance of a strong AU NH signal. most likely corresponding to the newly created base pair by the mutation in helix H3. Thus, once helix H3 is stabilized by replacing unpaired nucleotides within the internal loop to create a perfect helix, the A-C substitution at position -160 has no effect on the structure.

The fourth significant feature in the 3D structure is a small internal loop in helix H5 (five nts) involving nucleotides 24-25 and 34-37 (Fig. 2C and 3D). This is the most flexible region of the RNA. Nucleotides A35 and G36 are transiently solvent exposed, leading to relatively poor convergence for the neighboring base pairs as well (Fig. 3D). However, the entire helix retains the same structure with both allelic variants, suggesting it is not likely to be important for differential regulation of the promoter activity.

## Discussion

E-cadherin acts as a tumor suppressor in many carcinomas^8,9^, and its reduced expression is a hallmark of the epithelial-to-mesenchymal transition^10,11^. One mechanism of regulation of E-cadherin expression involves the recruitment of histone-modifying enzymes to the CDH1 promoter, which is regulated by a sense promoter-associated ncRNA transcribed independently of the main promoter^16^. As illustrated schematically in Fig. 1A, the A-allele is linked to decreased transcriptional activity and silencing of the E-cadherin gene requires formation of a microRNA (miRNA)-guided Argonaute 1 complex on the sense paRNA, which then recruits the SUV39H1 methyltransferase to affect chromatin modifications, thereby reducing E-cadherin expression and increasing cancer risk. Within this non-coding RNA, an A/C SNP at position -160 from the CDH1 transcription start site is associated with significantly increased prostate cancer risk as a result of increased recruitment of epigenetic enzymes and reduced activity of the CDH1 promoter^16^. However, the SNP is distant from the isomiR-binding site in primary sequence and secondary structure, raising the question of how this single nucleotide change affects transcription. A possibility is that the effect of the SNP occurs at the three-dimensional structure level, yet the role of RNA structure in noncoding RNA function has been suggested much more often than it has been demonstrated, with some notable exceptions^32-34^.

In order to address the role of RNA structure in the regulation of E-cadherin, we determined the three-dimensional NMR structure of the 90 nucleotides domain within the paRNA responsible for recruitment of the isomiR and epigenetic enzymes, by using a divide and conquer approach. We show here that the paRNA folds into a well-defined three-dimensional structure, and that the structures of A and C variant SNPs differ significantly. This conformational change extends from the three-way junction where the SNP is located to reorganize helix H3, propagating the effect of the single nucleotide change to the larger three-way junction from which the isomer binding site emanates^16^. The conformational rearrangement is initiated by the disruption of the structure of the SNP three-way junction as a result of the AA to AC change (Fig. 4); however, the presence of a second distortion nearby, the asymmetric three nucleotides internal loop in helix H3 which is only two base pairs away from the junction, facilitates reorganization of the entire helix. This conformational change is facilitated by local conformational flexibility observed for both the three-way junction and the internal loop (Fig. 4). In fact, when the helix is stabilized by mutating the internal loop into a perfectly paired helix, the NMR spectra of A- and C-alleles are very similar, implying very similar structures (Fig. S19). Thus, internal loop facilitates reorganization of the 3-way junction and helix H3.

**Fig. 4.**
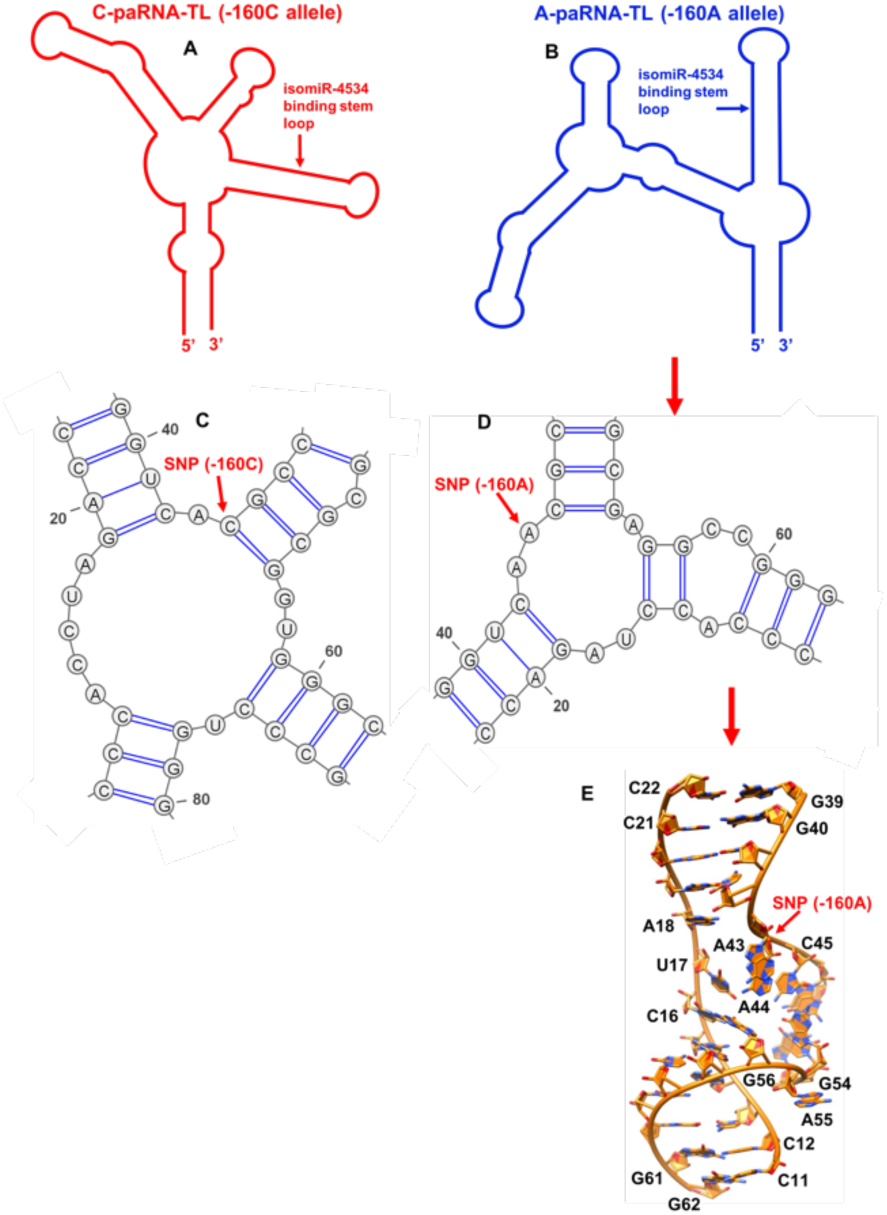
Schematic diagram showing the impact of SNP rs16260 on the structure of the CDH1 paRNA. A) and B) are cartoon representation of the secondary structures derived by NMR and SHAPE of the C- and A-paRNAs; C) and D) are close up view of the of the local secondary structure rearrangement induced by the C to A mutation at position - 160; E) close up view of the 3D structure of A-paRNA-TL corresponding to the secondary structure of panel D.

Through this structural mechanism, although the SNP is located approximately 50 nucleotides away from the isomiR-4534 binding site, the conformational rearrangement of helix H3 communicates the single nucleotide change to the entire paRNA domain, generating a plausible way to affect recruitment of factors that modulate E-cadherin expression. Our NMR structural studies provide a mechanistic basis for the effect of a single nucleotide change located away from the functional site and demonstrate that even minimal changes in sequence can significantly affect the function of ncRNAs by modulating RNA structure (Fig. 4). These results highlight a new mechanism by which polymorphic sites and somatic mutations in noncoding regions can affect the epigenetic landscape, inviting consideration of therapeutic targeting with small molecules. Several efforts have been made to discover compounds that impact cellular E-cadherin levels, but the molecular mechanism of action remains unclear^35,36^. Our NMR structural studies represent a first step towards the identification of small molecules that could restore E-cadherin expression in epithelial cancers by regulating protein expression at the RNA level.

## Methods

### RNA synthesis, purification and NMR sample preparation

All RNA constructs were prepared by *in vitro* transcription using *in house* purified T7 RNA polymerase and synthetic DNA templates (purchased from Integrated DNA Technologies), generally as reported^37,38^. 2′-O-methyl groups were incorporated with the last two residues at the 5′ end of the templates to reduce the addition of untemplated nucleotides at the 5′-end. ′Ultramer′ oligonucleotides were used for RNAs longer than 75 nucleotides. Deuterated RNA samples were prepared in the same way using selective deuterated rNTPs (D-H5, H3′, H4′, H5′ and H5′′) and ^13^C/^15^N-labeled samples were prepared using labeled rNTPs (Cambridge Isotope Laboratories). The resulting RNAs were purified for NMR studies by gel electrophoresis, electroelution and extensive dialysis, according to standard methods^37,38^. RNA concentrations used for NMR experiments were 0.5-1 mM.

### NMR experiments

All experiments were recorded on Bruker Avance 800 or/and 600 MHz NMR spectrometers equipped with ^1^H/ ^13^C/^15^N triple resonance cryogenic probes. Before each experiment, samples were freshly annealed by quick cooling after heating to 90 °C. The 1D ^1^H-^1^H spectra in 95% H_2_O/5% D_2_O were recorded using the excitation sculpting pulse sequence^39^. 2D ^1^H-^1^H nuclear Overhauser effect NOESY spectra were recorded at both 15 °C and 25 °C. 2D ^1^H-^1^H NOESY and 2D [^1^H-^15^N] HSQC spectra were recorded in 95% H_2_O/5% D_2_O in 20 mM phosphate (pH 6.0) and 0.01 mM EDTA, at 25 °C and 15 °C. In addition, 2D ^1^H-^1^H NOESY, ^1^H-^1^H TOCSY (spin lock 60-80 ms), 2D [^1^H-^13^C] HSQC, ^1^H-^13^C edited 3D NOESY-HSQC spectra were recorded in 99.9 % D_2_O buffer at 25 °C. The NOESY spectra were recorded with various mixing times in the range of 100-300 ms to facilitate spectral assignments and quantification of cross peak intensities for structure determination by comparison with peaks corresponding to fixed distances (e.g. H5-H6, H1′-H3′). ^1^H chemical shifts were referenced relative to external sodium 2,2-dimethyl-2-silapentane-5 sulfonate (DSS).

### NMR spectral assignments

In order to determine the structure of the A-paRNA domain including the A-allele, we used a divide-and-conquer approach and divided the RNA into four segments (Fig. S3), which overlap to generate the complete structure. We observed many NH peaks in the 1D NMR spectrum for the for the full paRNA domain and the shorter fragments revealing a well-folded structure (Fig. S5). Overlaying the 1D and 2D NOESY NMR spectra of the smaller segments on the spectra of the complete paRNA domain, revealed a highly transferable pattern of chemical shifts and NOESY cross-peaks for both the exchangeable and non-exchangeable protons (Figs S4-S5 and S11). In order to further simplify the spectra, we also prepared per-deuterated RNA samples (D-H5, H3′, H4′, H5′ and H5′′) for the full paRNA domain and the smaller fragments shown in Fig. S3, and recorded 2D-NOESY spectra at different mixing times as well. Overlaying the highly simplified, deuterated 2D NOESY spectra for the smaller domains on those of the full paRNA, we also observed considerable similarities in chemical shifts of the ribose sugar and aromatic region, to demonstrate the presence of highly transferable chemical shifts (Fig. S11). Thus, by collecting and comparing different 1D ^1^H, 2D ^1^H-^1^H NOESY and 3D NOESY-HSQC NMR spectra, we were able to transfer chemical shift assignments and NOEs from the smaller RNA domain to the complete paRNA domain (Figs S4-S5 and S11).

Spectral assignments of all fragments were facilitated by predicted RNA chemical shifts for helical regions^37,38^ and confirmed using established NOE helical “walk” patterns^37,38^ (Figs. S12-S15). Altogether, the analysis of multiple fragments enabled the bulk of the chemical shift assignments and NMR restraints to be obtained for the structural calculation of the complete A-paRNA domain.

### NMR structural constraints and structure calculations

In order to determine the 3D structure of the A-paRNA, we used the same divide and conquer approach used to assign chemical shifts because it allowed us to obtain a much larger number of distance restraints for structure calculation than would be possible if we just examined the complete RNA^17^. Its large molecular weight and asymmetric structure leads to rapid peak broadening due to fast relaxation of the NMR signal (Fig. S1). This approach is possible because RNA structure is highly modular, and individual stem-loops fold independently of each other; tertiary interactions, if present, are often weak and involve weaker long-range interactions between preformed RNA secondary structure motifs that only seldom disrupt the secondary structure^24,25^.

After dividing the A-paRNA domain into four partially overlapping smaller segments (Fig. S3) which, together, generate the complete A-paRNA structure, we obtained the bulk of distance restraints from NOE intensities used for the structure calculation by examining the smaller domains and by comparing them with the A-paRNA. This is possible because overlaying the 1D ^1^H and 2D ^1^H-^1^H NOESY NMR spectra of the smaller segments on the full A-paRNA spectra revealed a highly transferable pattern of chemical shifts and NOESY cross-peaks for both the exchangeable and non-exchangeable protons (Figs S4-S5 and S11). This observation demonstrates that the structure observed in the individual fragments is retained in the complete paRNA domain, because chemical shifts are very sensitive to even minor structural changes.

For structure calculations, NOE intensities were binned into ‘strong’ (2.5Å ± 0.7Å), ‘medium’ (3.5Å ± 1.5Å), and ‘weak’ (4.5Å ± 2.0Å) based on peak intensities relative to fixed A-form helical distances (e.g. H5-H6 = 2.5Å, H3′-H6/H8 = 3.5Å). Hydrogen bonds, planarity and dihedral restraints were included for base-paired nucleotides conforming to A-form helical patterns.

Xplor-NIH^27^ was used for structure calculations based on a simulated annealing protocol. The target function for refinement included NOE and hydrogen bond distance constraints (force constant = 50 kcal mol^−1^Å^2^) as well as dihedral torsion angle restraints (force constant = 200 kcal mol^−1^ rad^−2^). The simulated annealing procedure started from randomized coordinates and initially underwent high temperature torsion angle dynamics (8,000 steps at 3,500 K). At first, only NOE and Van Der Waals terms were active to fold the structure. Initial convergence was established when no NOE violations greater than 0.5 Å remained in the calculated structures. After this initial folding step, base-pair planarity and hydrogen-bonding restraints were examined for unambiguously established base pairs as identified from 2D ^1^H-^1^H NOESY and ^15^N-^1^H HSQC experiments. The bath temperature was gradually cooled from 3,500 to 298 K while introducing the van der Waals terms and incrementally raising the force constants for a number of terms (bond angles, impropers, dihedral angles, NOEs, van der Waals repulsion and ‘RAMA’). Following the final cooling step, the molecules underwent two sequential final Powell minimizations, first in torsion angle then in Cartesian space. The calculations were repeated multiple times and the lowest energy structure without distance (>0.5Å) or torsion angle violations (>5°) were used for further refinement steps incorporating a statistical base-base position potential for base-paired nucleotides. The 10 structures minimum energy structural ensemble was derived from 150 independent calculations using restraints from NOESY experiments and predicted values for A-form helical base-pairs (backbone and ribose dihedral angles, hydrogen-bond and planarity distance restraints).

A summary of the NMR restraints used in our structure calculation is provided in Table 1. Convergence was established when we observed no NOE violation greater than 0.5 Å or dihedral angle violations greater than 5° for the majority of structures within the final ensemble. The 3D structures were visualized with PyMol^40^ or Chimera^41^ and structural quality was analyzed using Molprobity^42^.

## Author contribution

Conceptualization, methodology, investigation, writing, reviewing and editing; graphics: SS Experimental design, writing, reviewing and editing: GV

## Acknowledgements

We are grateful to all members of the Varani groups for useful discussions and advice.

## References

1. Takeichi, M. et al. Cadherin cell adhesion receptors as a morphogenic regulator. Science. 251, 1451–1455 (1991).

2. van Roy, F. & Berx, G. The cell–cell adhesion molecule E-cadherin. Cell. Mol. Life Sci. 65, 3756–3788 (2008).

3. Oda, H. & Takeichi, M. Evolution: structural and functional diversity of cadherin at the adherens junction. J. Cell Biol. 193, 1137–1146 (2011).

4. Richards, F.M., McKee, S.A., Rajpar, M.H., Cole, T.R., Evans, D.G., Jankowski, J.A., McKeown, C., Sanders, D.S. & Maher, E.R. Germline E-cadherin gene (CDH1) mutations predispose to familial gastric cancer and colorectal cancer. Hum Mol Genet. 8, 607–610 (1999).

5. Hajra, K.M. & Fearon, E.R. Cadherin and catenin alterations in human cancer. GENES, CHROMOSOMES & CANCER. 34, 255–268 (2002).

6. Ozawa, M., Baribault, H. & Kemler, R. The cytoplasmic domain of the cell adhesion molecule uvomorulin associates with three independent proteins structurally related in different species. EMBO J. 8, 1711–1717 (1989).

7. Kim, N.G., Koh, E., Chen, X. & Gumbiner, B.M. E-cadherin mediates contact inhibition of proliferation through Hippo signaling-pathway components. Proc Natl Acad Sci USA. 108, 11930–11935 (2011).

8. Bray, F., Ferlay, J., Soerjomataram, I., Siegel, R.L., Torre, L.A. & Jemal, A. Global cancer statistics 2018: GLOBOCAN estimates of incidence and mortality worldwide for 36 cancers in 185 countries. CA Cancer J Clin. 68, 394–424 (2018).

9. Pan, R., Zhu, M., Yu, C., Lv, J., Guo, Y., Bian, Z., Yang, L., Chen, Y., Hu, Z., Chen, Z., Li, L., Shen, H. China Ka-doorie Biobank Collaborative Group, Cancer incidence and mortality: a cohort study in China, 2008-2013. Int J Cancer. 141, 1315–1323 (2017).

10. Yang, J. & Weinberg, R.A. Epithelial-mesenchymal transition: at the crossroads of development and tumor metastasis. Dev Cell. 14, 818–829 (2008).

11. Lamouille, S., Xu, J. & Derynck, R. Molecular mechanisms of epithelial-mesenchymal transition. Nat Rev Mol Cell Biol. 15, 178–196 (2014).

12. Cano, A., Pérez-Moreno, M. A., Rodrigo, I., Locascio, A., Blanco, M. J., del Barrio, M.G., Portillo, F. & Nieto, M. A. The transcription factor snail controls epithelial-mesenchymal transitions by repressing E-cadherin expression. Nat. Cell Biol. 2, 76–83 (2000).

13. Graff, J.R., Herman, J.G., Lapidus, R.G., Chopra, H., Xu, R., Jarrard, D.F., Isaacs, W.B., Pitha, P.M., Davidson, N.E. & Baylin, S.B. E-cadherin expression is silenced by DNA hypermethylation in human breast and prostate carcinomas. Cancer Res. 55, 5195–5199 (1995).

14. Yoshiura, K., Kanai, Y., Ochiai, A., Shimoyama, Y., Sugimura, T. & Hirohashi, S. Silencing of the E-cadherin invasion-suppressor gene by CpG methylation in human cancers. Proc Natl Acad Sci USA. 92, 7416 –7419 (1995).

15. Li, L.C., Chui, R.M., Sasaki, M., Nakajima, K., Perinchery, G., Au, H. C., Nojima, D., Carroll, P. & Dahiya, R. A Single Nucleotide Polymorphism in the E-cadherin Gene Promoter Alters Transcriptional Activities. Cancer Res. 60, 873–876 (2000).

16. Pisignano, G., Napoli, S., Magistri, M., Mapelli, S.N., Pastori, C., Marco, S.D., Civenni, G., Albino, D., Enriquez, C., Allegrini, S., Mitra, A., D’Ambrosio, G., Mello-Grand, M., Chiorino, G., Garcia-Escudero, R., Varani, G., Carbone, G.M. & Catapano, C.V. A promoter-proximal transcript targeted by genetic polymorphism controls E-cadherin silencing in human cancers. Nat Commun. 8, 15622 (2017).

17. Barnwal, R.P., Loh, E., Godin, K.S., Yip, J., Lavender, H., Tang, C.M. & Varani, G. Structure and mechanism of a molecular rheostat an RNA thermometer that modulates immune evasion by Neisseria meningitidis. Nucleic Acids Research. 44, 9426–9437 (2016).

18. Walker, M.J., Shortridge, M. D., Albin, D. D., Cominsky, L.Y. & Varani, G. Structure of the RNA Specialized Translation Initiation Element that Recruits eIF3 to the 5′-UTR of c-Jun. J. Mol. Biol. 432, 1841–1855 (2020).

19. Lawrence, D.C., Stover, C.C., Noznitsky Wu, J. Z. & Summers, M.F. Structure of the Intact Stem and Bulge of HIV-1 ψ-RNA Stem-Loop SL1. J. Mol. Biol. 326, 529–542 (2003).

20. Keane, S. C. & Summers, M.F. NMR Studies of the Structure and Function of the HIV-1 5’-Leader, Viruses. 8, 338 (2016).

21. Duss, O., Michel, E., Yulikov, M., Schubert, M., Jeschke, G. & Allain, F.H.T. Structural basis of the non-coding RNA RsmZ acting as a protein sponge. Nature. 509, 588–592 (2014).

22. Duss, O., Lukavsky, P.J. & Allain, F.H.T. Isotope labeling and segmental labeling of larger RNAs for NMR structural studies. Advances in Experimental Medicine and Biology. 992, 121–144 (2012).

23. Sharma, S. & Varani, G. NMR structure of Dengue West Nile viruses stem-loop B: A key cis-acting element for flavivirus replication. Biochem. Biophys. Res. Commun. 531, 522–527 (2020).

24. Wu, M. & Tinoco Jr. I. RNA folding causes secondary structure rearrangement. Proc. Natl. Acad. Sci. USA. 95 11555–11560 (1998).

25. Celander, D.W. & Cech, T.R. Visualizing the higher order folding of a catalytic RNA molecule. Science. 251, 401–407(1991).

26. Tinoco Jr. I. & Bustamante, C. How RNA Folds, J. Mol. Biol. 293, 271–281(1999).

27. Schwieters, C.D., Kuszewski, J.J., Tjandra, N. & Clore, G.M. The Xplor-NIH NMR molecular structure determination package, J. Magn. Reson. 160, 65–73 (2003).

28. Lescoute, A. & Westhof, E. Topology of three-way junctions in folded RNAs. RNA. 12 83–93 (2006).

29. Laing, C. & Schlick, T. Analysis of four-way junctions in RNA structures, J. Mol. Bio. 390, 547–559 (2009).

30. Gutell, R.R., Cannone, J.J., Shang, Z. Du, Y. & Serra, M.J. A story: unpaired adenosine bases in ribosomal RNAs. J. Mol. Bio. 304, 335–354 (2000).

31. Gutell, R.R., Weiser, B., Woese, C.R. & Noller, H.F. Comparative anatomy of 16S-like ribosomal RNA. Prog. Nucl. Acid Res. Mol. Biol. 32, 155–216 (1985).

32. Jones, A.N. & Sattler, M. Challenges and perspectives for structural biology of lncRNAs-the example of the Xist lncRNA A-repeats. Journal of Molecular Cell Biology. 11, 845–859 (2019).

33. Pintacuda, G., Young, A.N. & Cerase, A. Function by Structure: Spotlights on Xist Long Non-coding RNA. Front. Mol. Biosci. 4, 90 (2017).

34. Brown, J.A., Bulkley, D., Wang, J., Valenstein, M.L., Yario, T. A., Steitz, T.A. & Steitz, J.A. Structural insights into the stabilization of MALAT1 noncoding RNA by a bipartite triple helix. Nat Struct Mol Biol. 21, 633–640 (2014).

35. Song, Y., Ye, M., Zhou, J., Wang, Z. & Zhu, X. Targeting E-cadherin expression with small molecules for digestive cancer treatment. Am J Transl Res. 11, 3932–3944 (2019).

36. Song, Y., Ye, M., Zhou, J., Wang, Z. & Zhu, X. Restoring E-cadherin Expression by Natural Compounds for Anticancer Therapies in Genital and Urinary Cancers. Mol Ther Oncolytics. 14, 130–138 (2019).

37. Varani, G., Aboulela, F. & Allain, F.H.T. NMR investigation of RNA structure. Prog. NMR Spectrosc. 29, 51–127 (1996)

38. Varani, G. & Tinoco Jr. I. RNA structure and NMR spectroscopy. Q. Rev. biophysics. 24, 479–532 (1991).

39. Hwang, T.L. & Shaka, A.J. Water suppression that works. Excitation sculpting using arbitrary waveforms and pulsed field gradients, J. Magn. Reson., Series A. 112, 275–279 (1995).

40. Schrodinger, L.L.C. The PyMOL Molecular Graphics System, 2010. Version 1.3r1.

41. Pettersen, E.F., Goddard, T.D. & Huang, C.C. UCSF Chimera-a visualization system for exploratory research and analysis. J. Comput. Chem. 25, 1605–1612 (2004).

42. Davis, I.W., Murray, L.W., Richardson, J.S. & Richardson, D.C. MolProbity: structure validation and all-atom contact analysis for nucleic acids and their complexes. Nucleic Acids Res. 32, 615–619 (2004).

