## supplementary file for "A functional SNP regulates E-cadherin expression by dynamically remodeling the 3D structure of a promoter-associated non-coding RNA transcript"

### Supporting Figures:

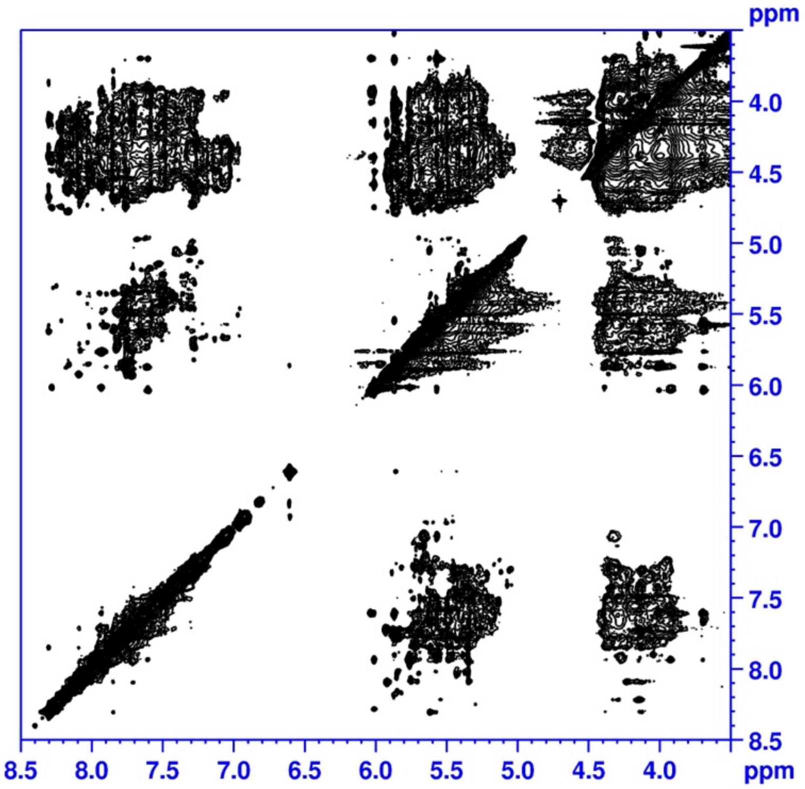


**Fig. S1.** 2D NOESY spectrum (non-exchangeable protons) for A-paRNA; the relatively poor quality of the spectra, as reflected in broad and highly overlapped signals for the non-exchangeable protons, is expected for an RNA of this size and would make structure determination impossible.

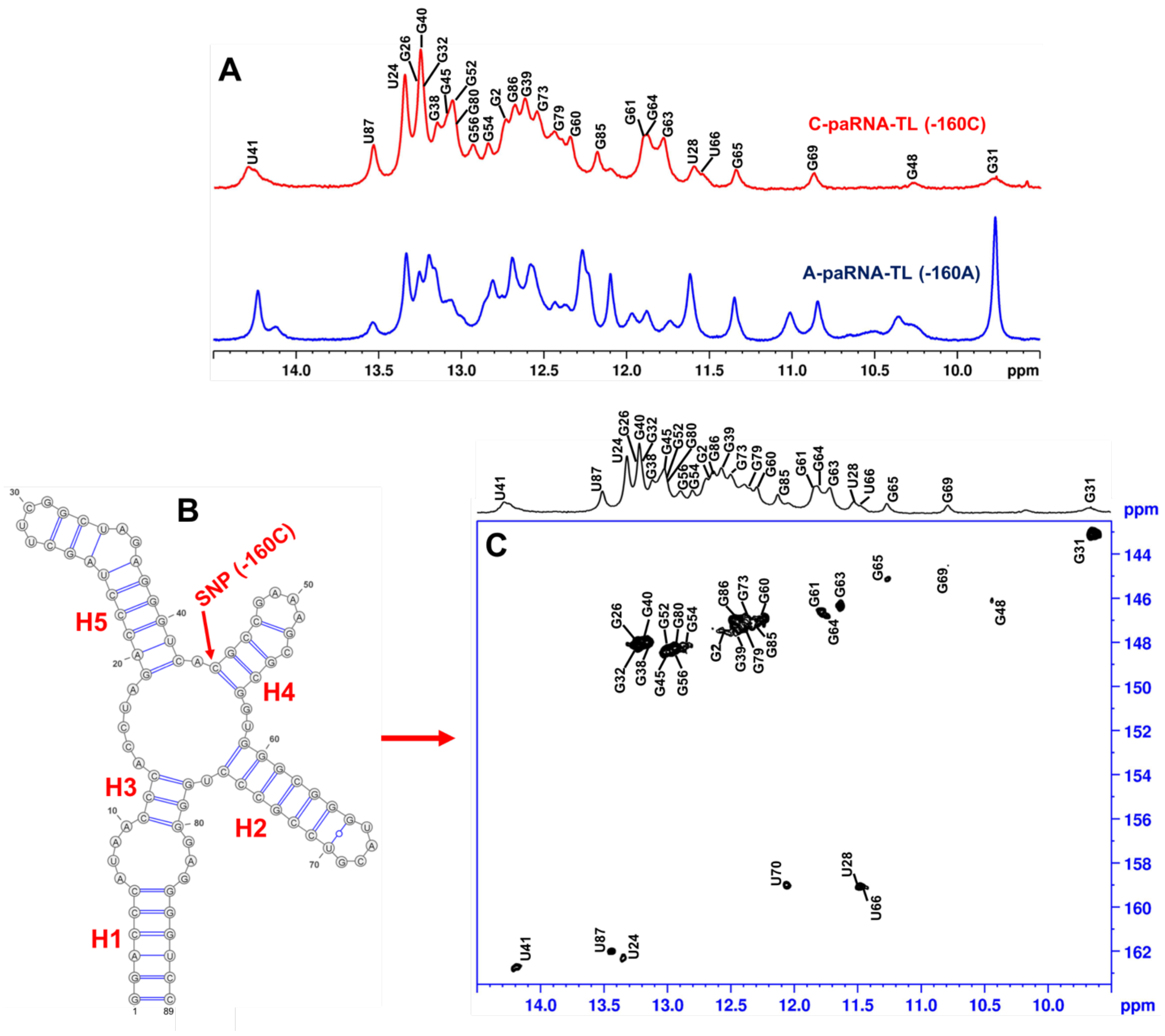


**Fig. S2.** A and C variants differ substantially in their secondary structure. An overlay of the imino $^{1}$H NMR spectra for the A-paRNA-TL (-160A allele) and C-paRNA-TL (-160C allele). B) Secondary structure of C-paRNA-TL, as established from the NMR assignments and fully consistent with the SHAPE analysis; C) 2D-$^{15}$N-HSQC spectra of the same RNA construct, with NMR assignments.

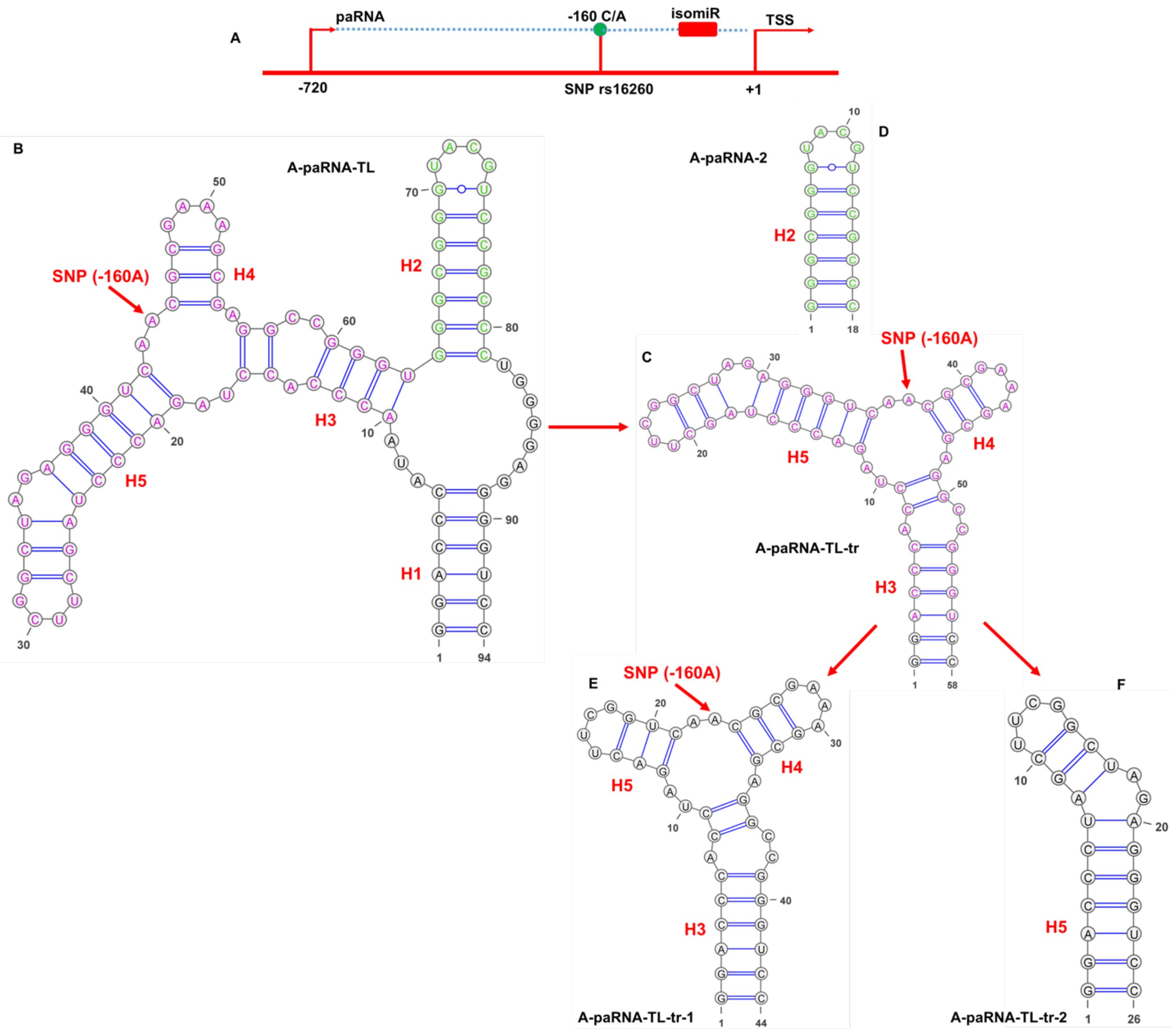


**Fig. S3**. A) Schematic diagram of the genomic region that immediately precedes the transcription start site of the CDH1 gene; the independent transcription start site for the paRNA; the SNP rs16260 (green circle) and the isomiR-4534 binding site (red rectangle) are marked; they are separated in both primary sequence and secondary structure; B) SHAPE-derived secondary structure (confirmed by NMR in this study) of the A-paRNA construct, which was divided into four segments which overlap to generate the complete structure; these five constructs were all used in the NMR studies.

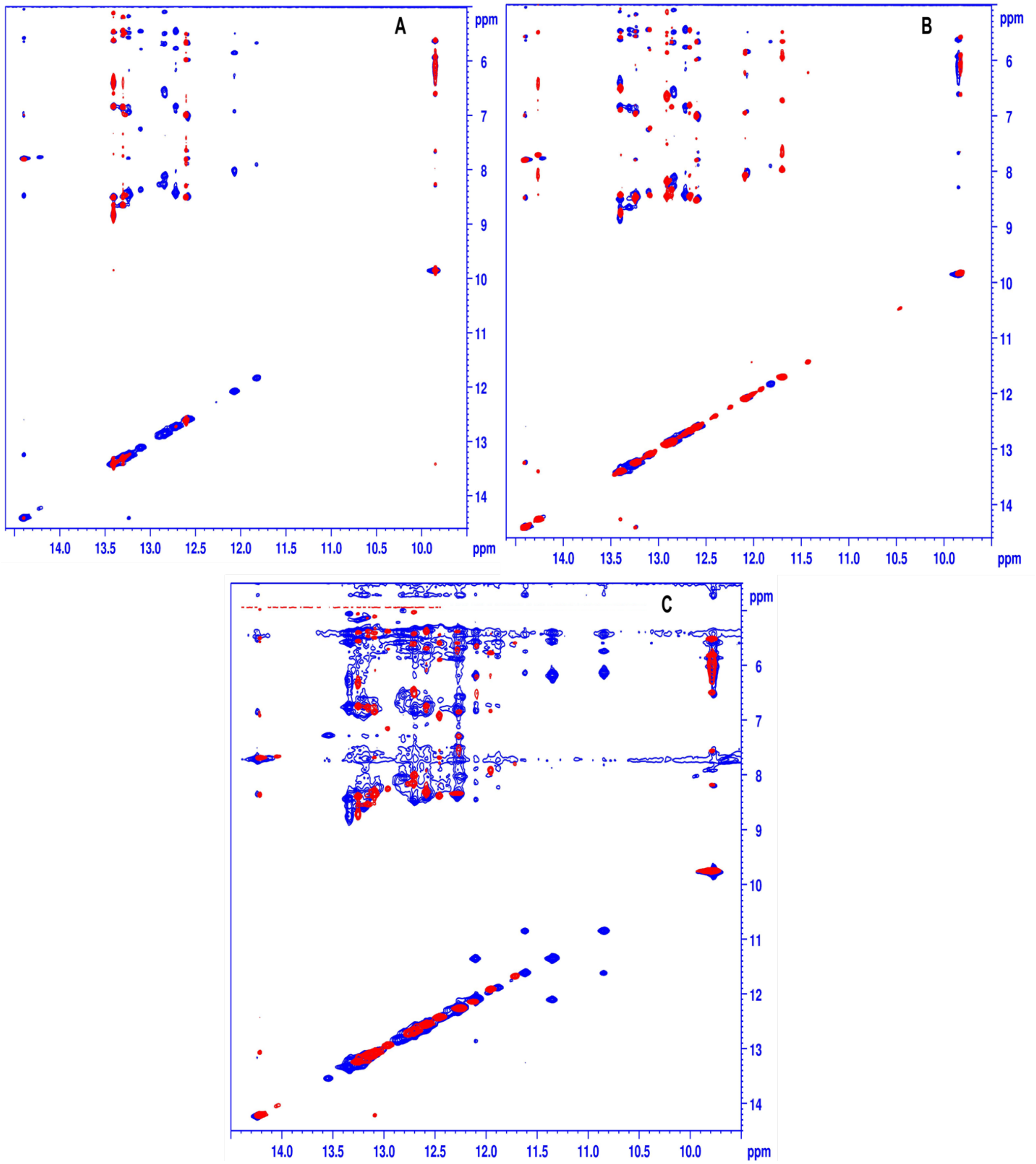


**Fig. S4.** Overlay of the imino regions of 2D NOESY NMR spectra. A) Overlay of A-paRNA-TL-tr (blue) and A-paRNA-TL-tr-2 (red), with reference to Fig. S3; B) A-paRNA-TL-tr (blue) and A-paRNA-TL-tr-1 (red); C) A-paRNA-TL (blue) and A-paRNA-TL-tr (red). The considerable similarities in the imino resonance peak positions (as in Figure S5) and pattern of NOE cross peaks demonstrate the presence of very similar secondary structures.

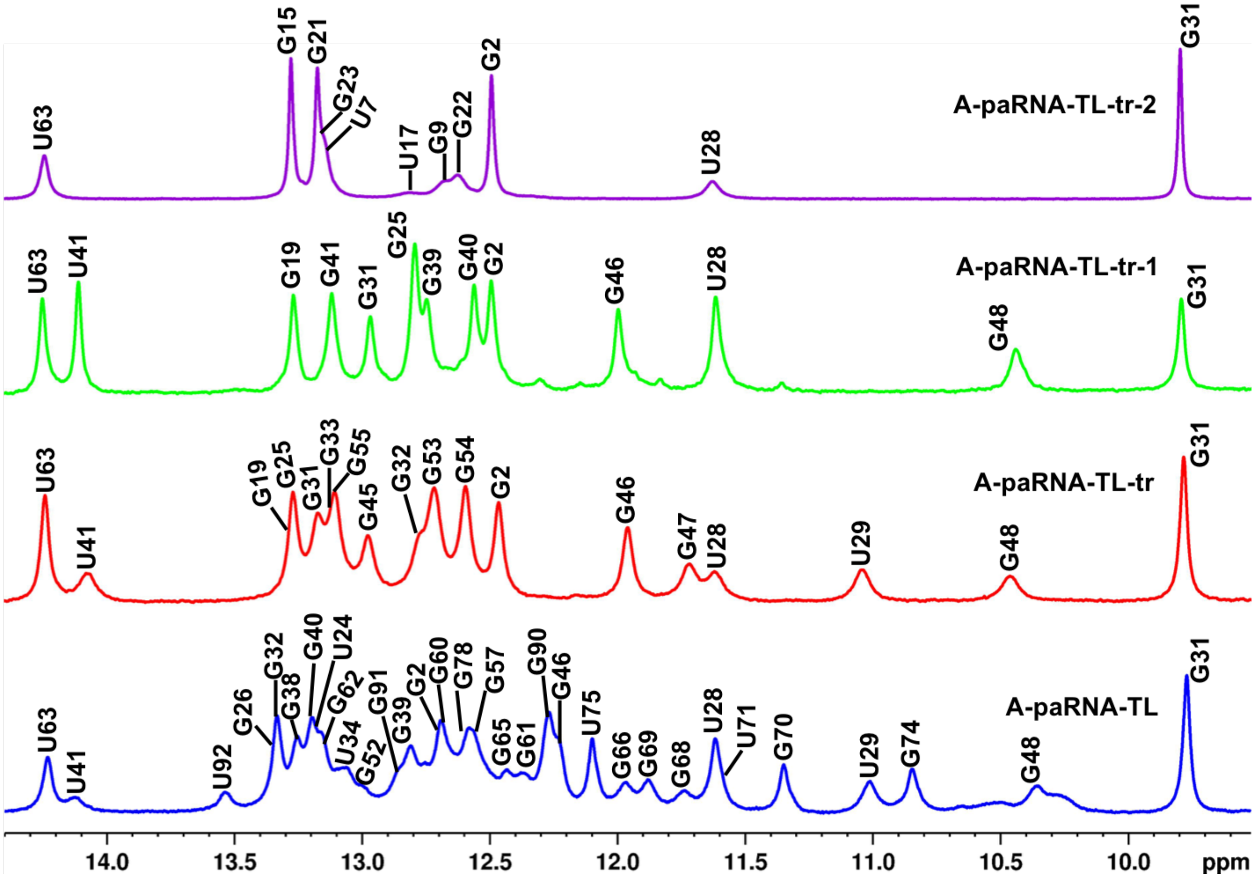


**Fig. S5**. Overlay of the 1D $^{1}$H NMR spectra (imino region only) for the A-paRNA-TL, A-paRNA-TL-tr, A-paRNA-TL-tr-1 and A-paRNA-TL-tr-2 constructs, with assignments; with reference to the secondary structures of Fig. S3. The remarkable similarities in the imino resonances demonstrate the presence of very similar secondary structures across all four RNAs. In other words, the secondary structure of the full A-paRNA is fully preserved in each of the individual fragments.

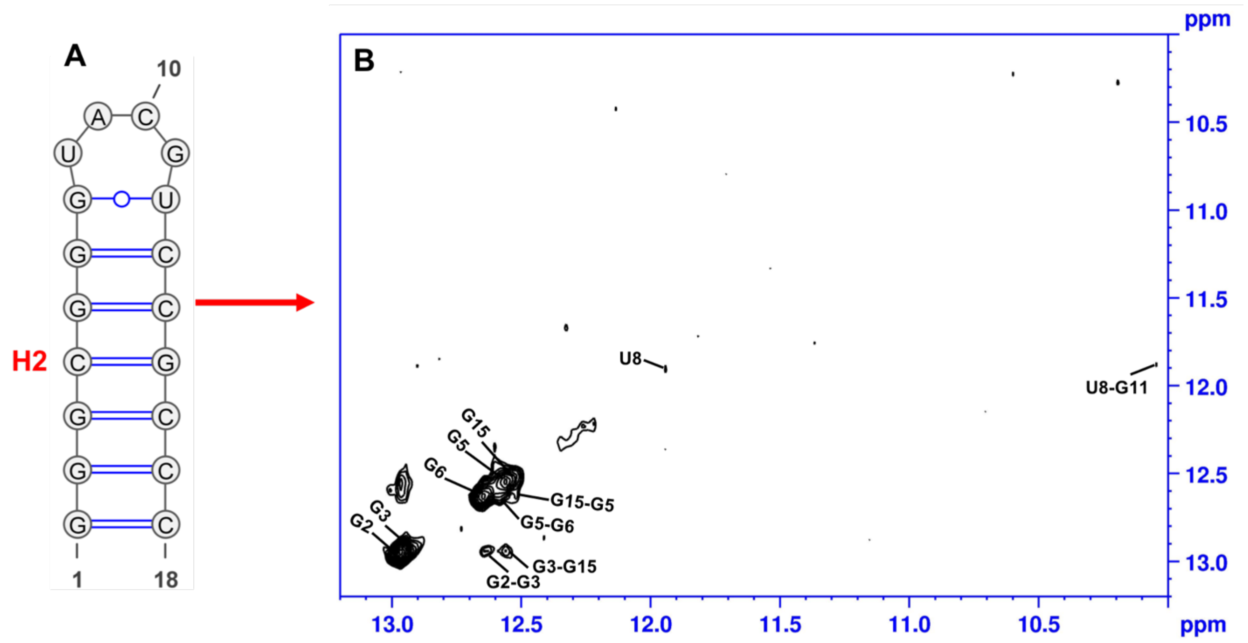


**Fig. S6**. A) NMR-derived secondary structure of A-paRNA-2, corresponding to the isomiR-binding site, helix H2; B) Imino region of the 2D-NOESY NMR spectrum of A-paRNA-2, with assignments that verify the secondary structure as shown.

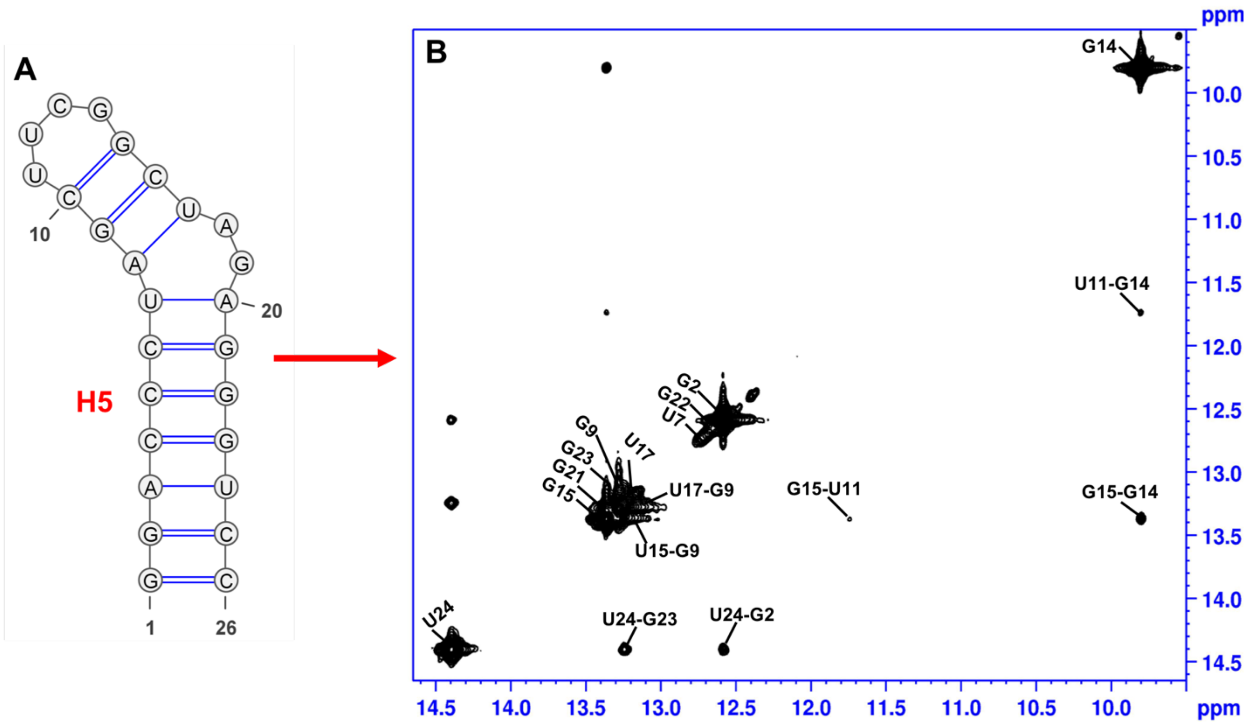


**Fig. S7**. A) NMR-derived secondary structure of A-paRNA-tr-2, corresponding to the large stem-loop emerging from the 3-way junction where the SNP is located, helix H5; B) Imino region of the 2D-NOESY NMR spectrum of A-paRNA-tr-2, with assignments that verify the secondary structure, as shown.

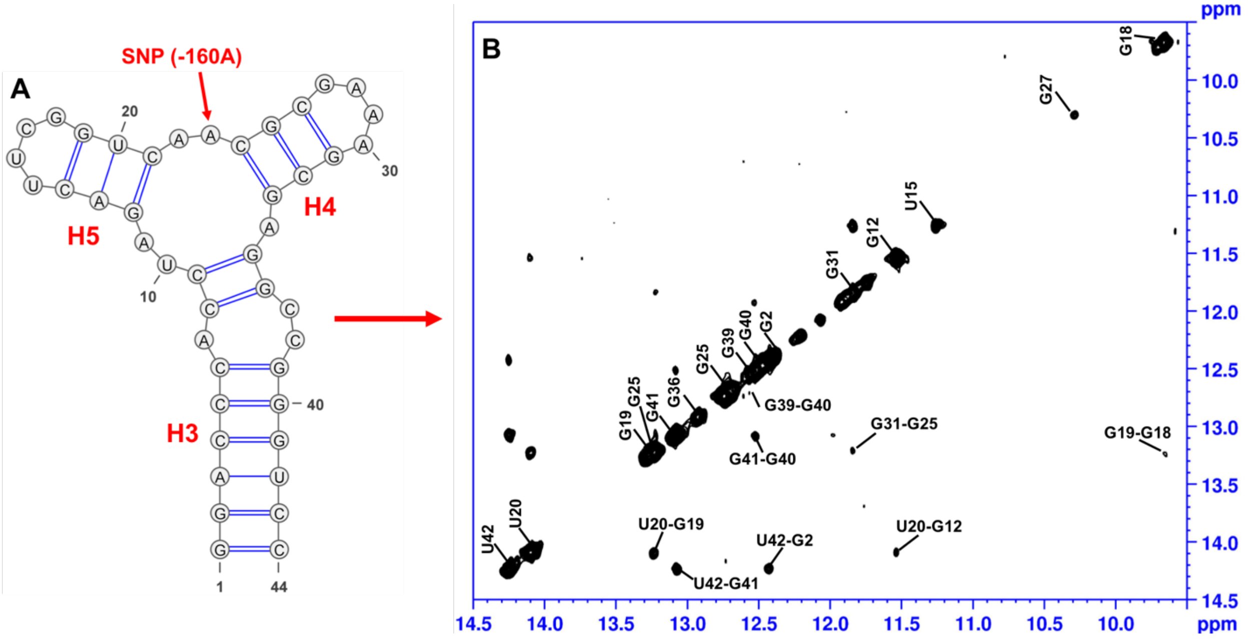


**Fig. S8**. A) NMR-derived secondary structure of A-paRNA-tr-1 which isolates the 3-way junction formed by helices H3, H4 and H5, where the SNP is located within a smaller RNA amenable to high-resolution investigation; B) Imino region of the 2D-NOESY NMR spectrum of A-paRNA-tr-1, with assignments that verify the secondary structure, as shown.

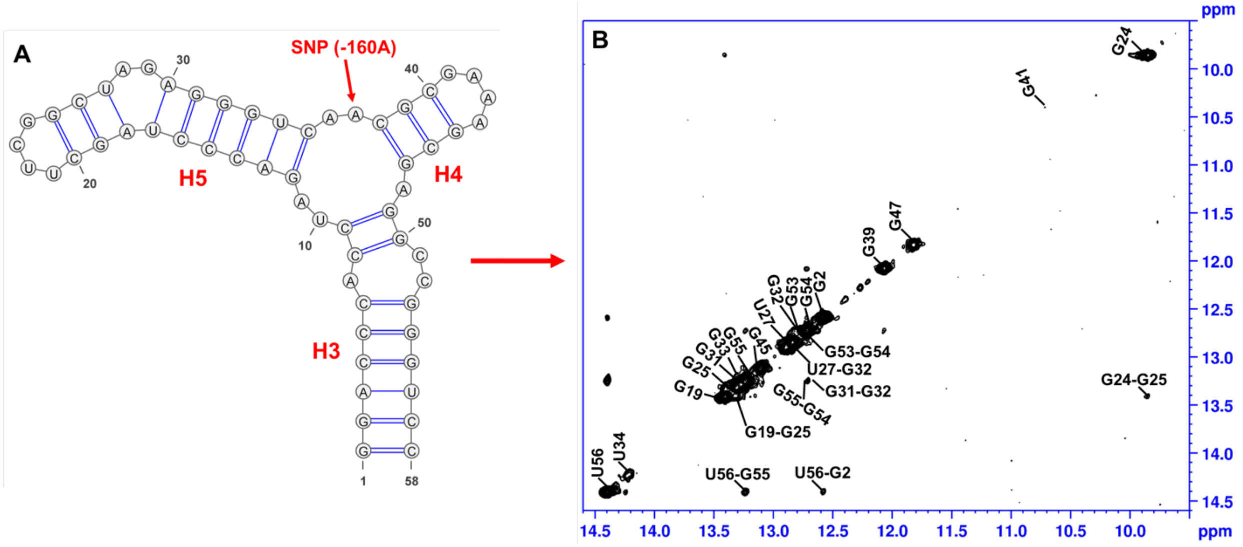


**Fig. S9**. A) NMR-derived secondary structure of A-paRNA-tr, that isolates the 3-way junction where the SNP is located; B) Imino region of the 2D-NOESY NMR spectrum of A-paRNA-tr, with assignments that verify the secondary structure, as shown.

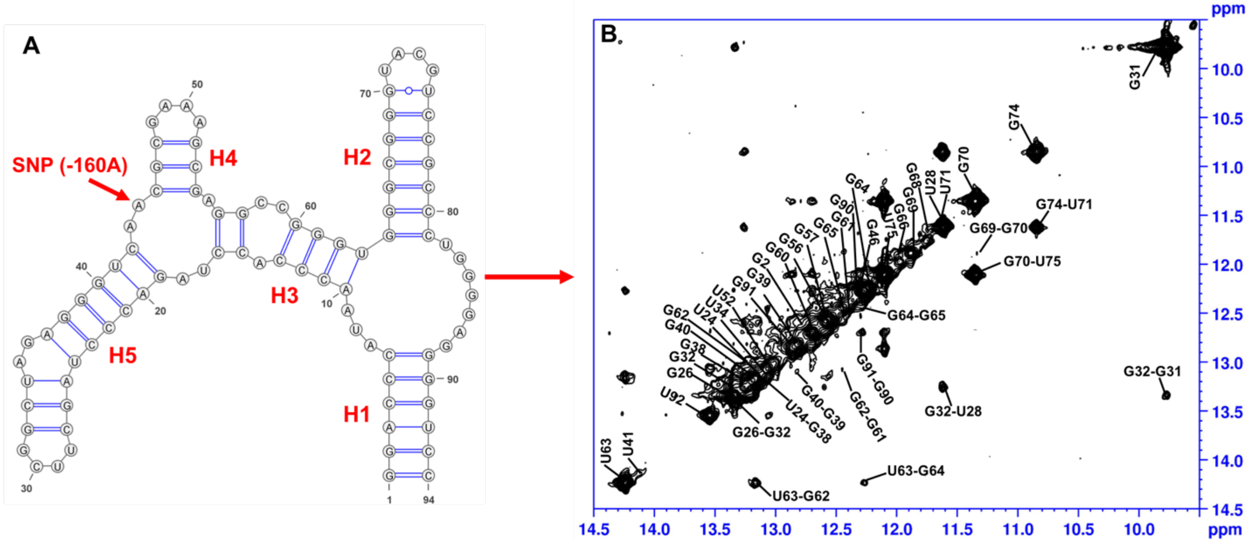

**Fig. S10**. A) SHAPE-derived secondary structure of A-paRNA-TL, with tetraloops that stabilize the structure and reduce aggregation; B) Imino region of the 2D-NOESY NMR spectrum of A-paRNA-TL, with assignments that verify the secondary structure, as shown, fully consistent with the SHAPE analysis.

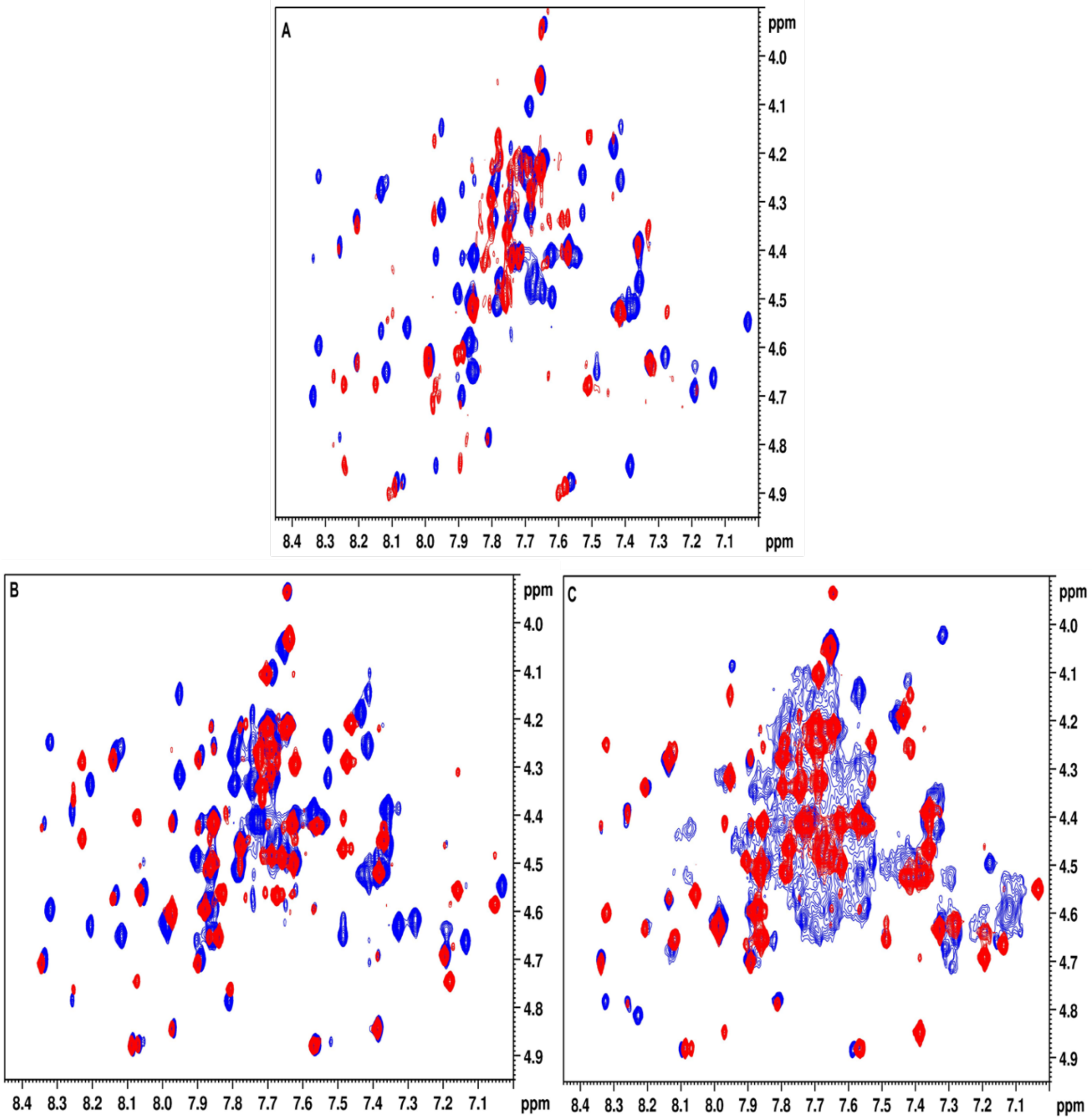


**Fig. S11**. Overlay of the $D_2O$ 2D NOESY spectra of perdeuterated RNAs reveals a highly transferable pattern of chemical shifts and NOESY cross peaks. A) Overlay of the H2′ to H6/H8 region of A-paRNA-TL-tr (blue) and A-paRNA-TL-tr-2 (red); B) the H2′ to H6/H8 region of A-paRNA-TL-tr (blue) and A-paRNA-TL-tr-1 (red); C) H2′ to H6/H8 region of A-paRNA-TL (blue) and A-paRNA-TL-tr (red); even with extensive deuteration, overlap is considerable in this last spectrum, and demonstrates the necessity of the divide-and-conquer approach.

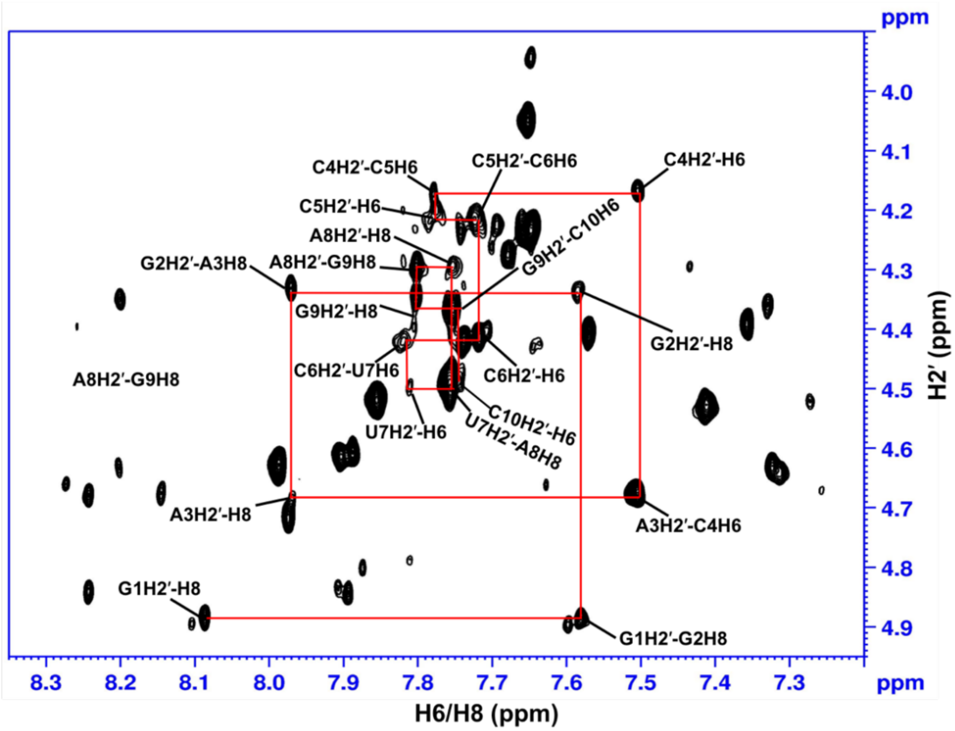


**Fig. S12**. The H2′ to H6/H8 'helical walk', annotated from G1 to C10, for A-paRNA-tr2, plotted from a 2D NOESY spectrum recorded at 25 °C with (H6/H8, H1′, H2′ but D3′, D4′, D5′/D5″ and D5) ribose deuteration.

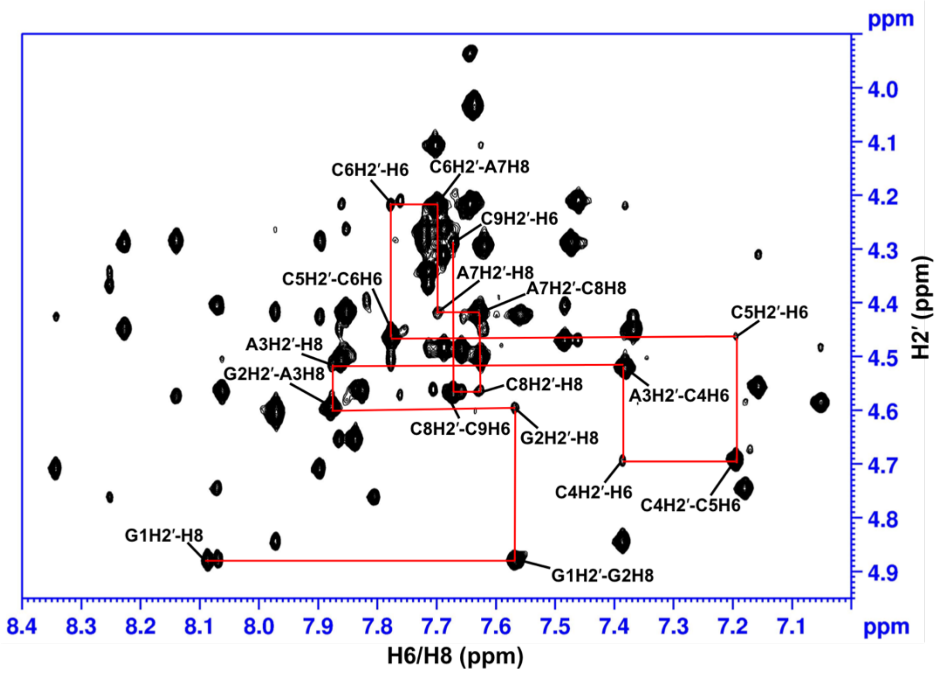


**Fig. S13.** The H2′ to H6/H8 'helical walk', annotated from G1 to C9, for A-paRNA-tr1, plotted from a 2D NOESY spectrum recorded at 25 °C with (H6/H8, H1′, H2′ but D3′, D4′, D5′/D5″ and D5) ribose deuteration.

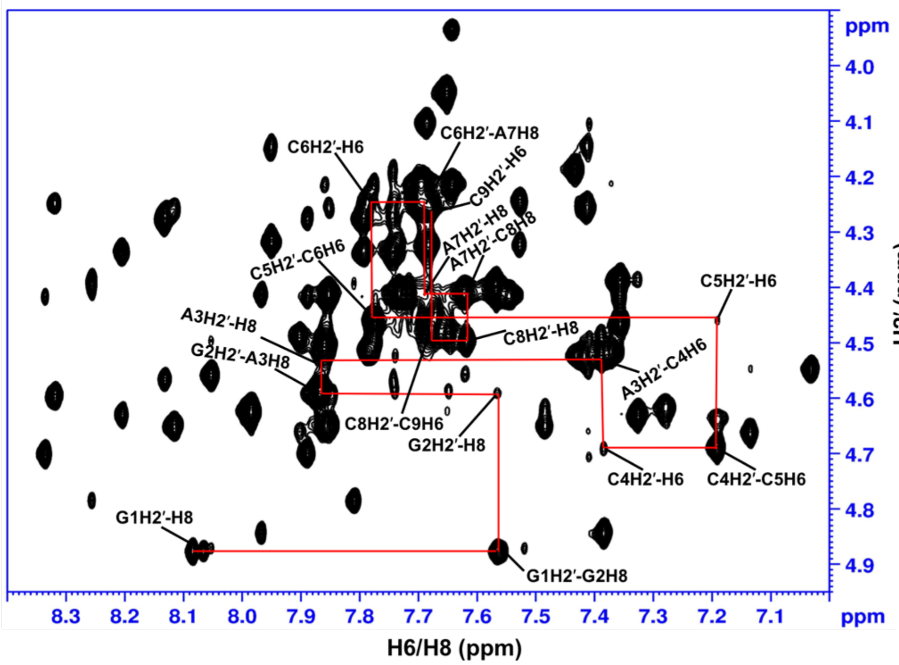


**Fig. S14.** The H2′ to H6/H8 'helical walk', annotated from G1 to C9, for A-paRNA-tr, plotted from the 2D NOESY spectrum recorded at 25 °C with (H6/H8, H1′, H2′ but D3′, D4′, D5′/D5″ and D5) ribose deuteration.

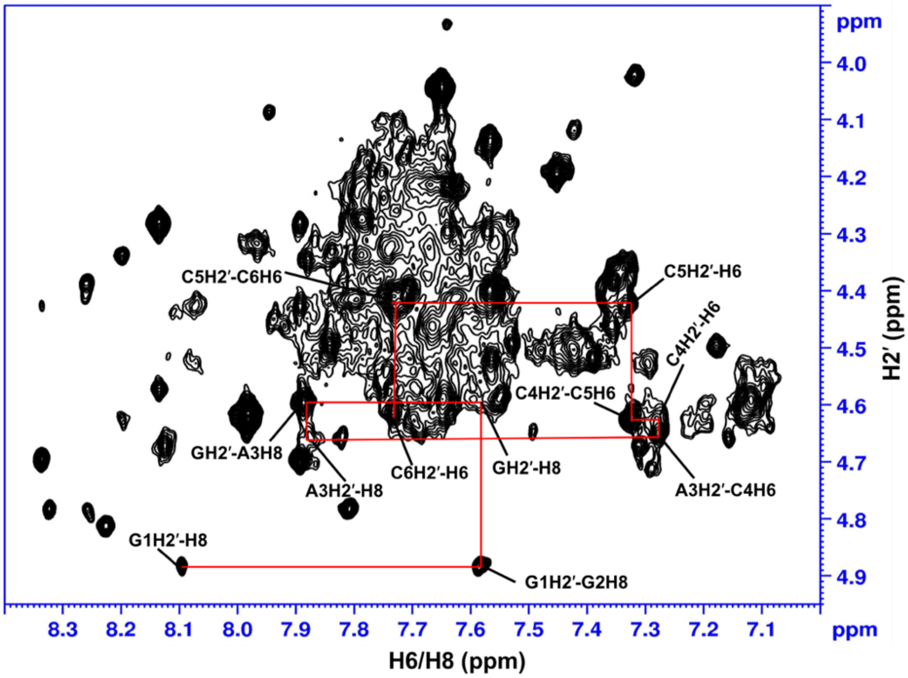


**Fig. S15.** The H2′ to H6/H8 'helical walk', annotated from G1 to C6, for A-paRNA-TL, plotted from the 2D NOESY spectrum recorded at 25 °C with (H6/H8, H1′, H2′ but D3′, D4′, D5′/D5″ and D5) ribose deuteration; even with extensive deuteration, overlap is considerable and demonstrates the necessity of the divide-and-conquer approach.

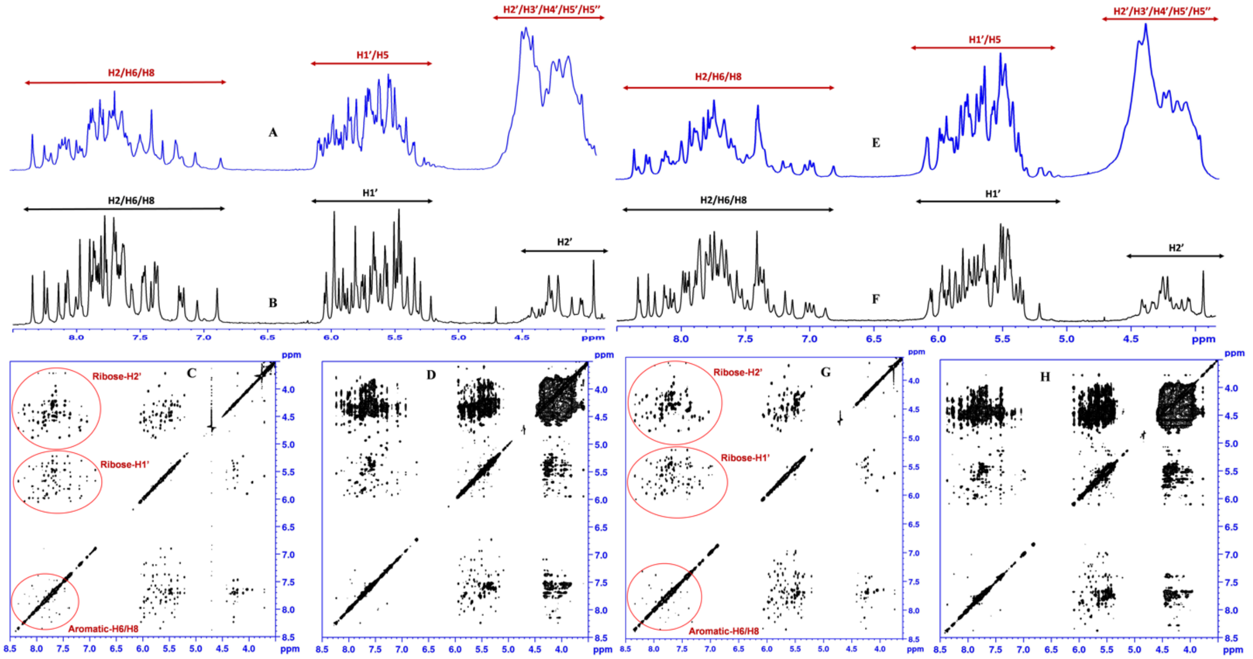


**Figure S16.** Base and ribose deuteration greatly improve the quality of the NMR spectra of these relatively large RNAs; 1D $^{1}$H NMR spectra collected without A) and with B) (H6/H8, H1′, H2′, D3′, D4′, D5′/D5″and D5-ribose deuteration; C) and D) are 2D NOESY spectra recorded at 25 °C for the A-paRNA-tr-1. E) 1D $^{1}$H NMR spectra with B) and without F) (H6/H8, H1′, H2′, D3′, D4′, D5′/D5″and D5-ribose deuteration); NOESY spectra with G) and without H) selective deuteration at 25 °C for the A-paRNA-tr.

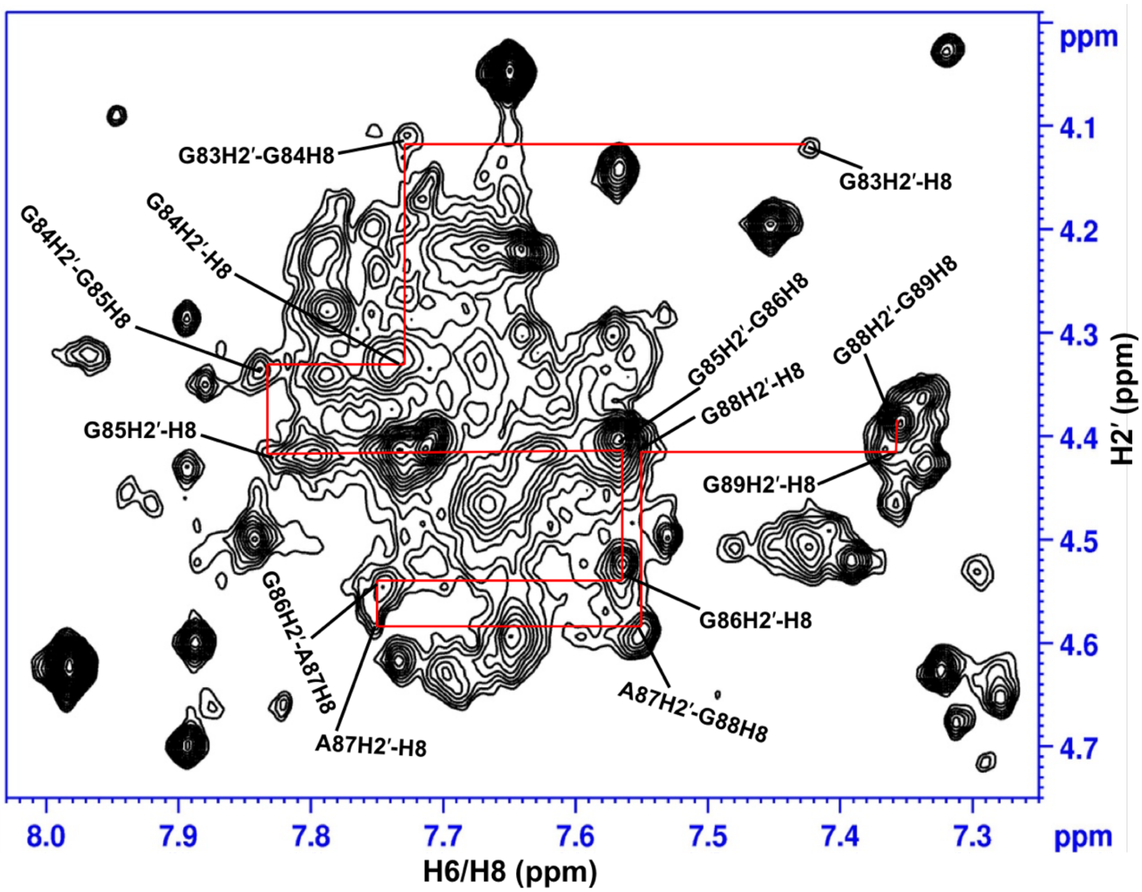


**Fig. S17.** The H2′ to H6/H8 'helical walk', labeled from G83 to G89, for A-paRNA-TL. The 2D NOESY spectrum was recorded at 25 °C with (H6/H8, H1′, H2′ but D3′, D4′, D5′/D5″ and D5) ribose deuteration.

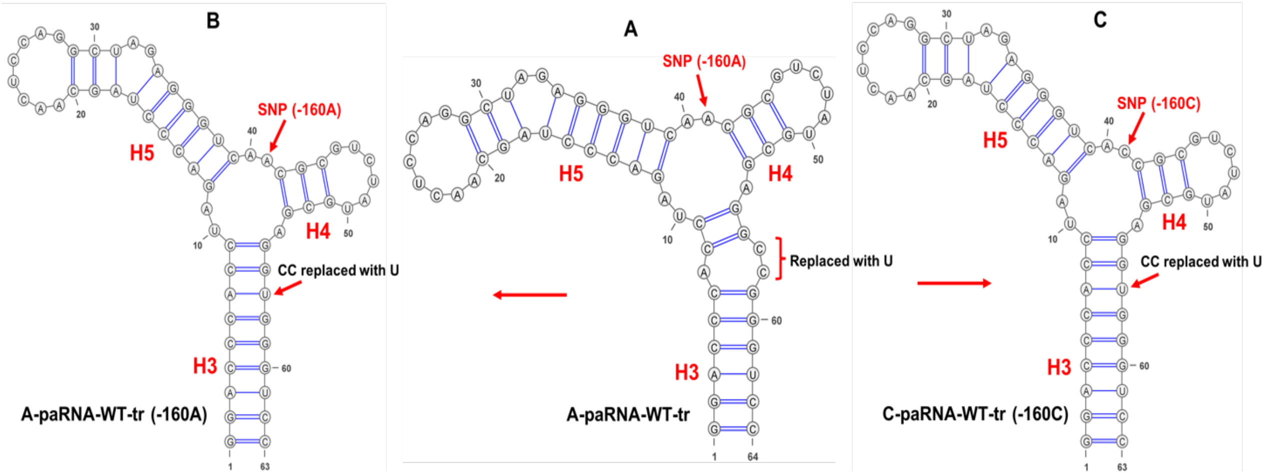


**Fig. S18.** A) Secondary structure of A-paRNA-TL-tr; B) and C) are the secondary structures of A and C-variants of the same RNAs, but in these two RNAs, C58 and C59, which are part of the internal loop in helix H3, are replaced with U, as shown with red arrows, to generate a perfectly paired helix H3. These constructs contain the natural loop sequences and not stabilizing tetraloops.

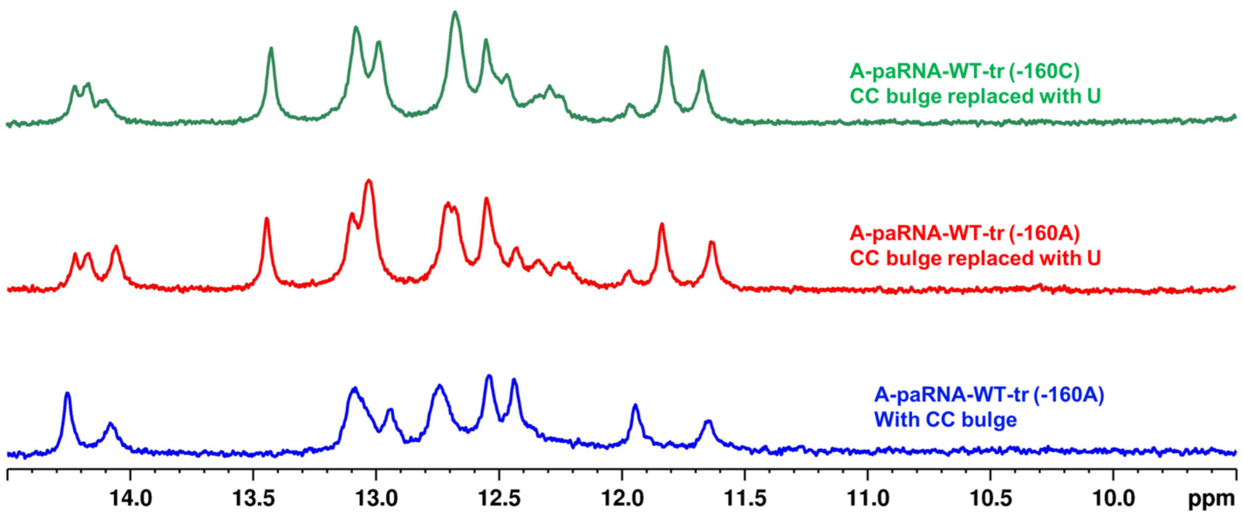


**Fig. S19.** Comparison of the 1D imino $^{1}H$ NMR spectra for the three RNAs of Fig. S18, corresponding to the helix H3, H4 and H5 portion of the paRNA with the A allele (bottom spectrum); the same RNA, but with C58C59 mutated to U to eliminate the internal loop and create a perfectly paired helix H3 (center) and, finally, the same perfectly paired helix H3, but with the C allele at position -160 in the three-way junction. The three spectra are very similar, with the exception of the sharp resonance at about 13.5 ppm, most likely corresponding to the new AU base pair created by the mutation. Once helix H3 is stabilized by replacing the internal loop with a perfectly paired helix, the A to C substitution in the 3-way junction does not induce a conformational change.
